# Genomics of altitude-associated wing shape in two tropical butterflies

**DOI:** 10.1101/2020.12.05.412882

**Authors:** Gabriela Montejo-Kovacevich, Patricio A. Salazar, Sophie H. Smith, Kimberly Gavilanes, Caroline N. Bacquet, Yingguang Frank Chan, Chris D. Jiggins, Joana I. Meier, Nicola J. Nadeau

**Affiliations:** Department of Zoology, University of Cambridge, Cambridge, CB2 3EJ, UK; Animal and Plant Sciences, University of Sheffield, Sheffield, S9 2TN, UK; Universidad Regional Amazónica de Ikiam, Tena, Ecuador; Friedrich Miescher Laboratory of the Max Planck Society, Max Planck Ring 9, 72076 Tübingen, Germany; St John’s College, University of Cambridge, Cambridge, CB2 3EJ, UK

## Abstract

Understanding how organisms adapt to their local environment is central to evolution. With new whole-genome sequencing technologies and the explosion of data, deciphering the genomic basis of complex traits that are ecologically relevant is becoming increasingly feasible. Here we study the genomic basis of wing shape in two Neotropical butterflies that inhabit large geographical ranges. *Heliconius* butterflies at high elevations have been shown to generally have rounder wings than those in the lowlands. We reared over 1100 butterflies from 71 broods of *H. erato* and *H. melpomene* in common-garden conditions and show that wing aspect ratio, i.e. elongatedness, is highly heritable in both species and elevation-associated wing shape differences are maintained. Genome-wide associations with a published dataset of 666 whole genomes from across a hybrid zone, uncovered a highly polygenic basis to wing shape variation in the wild. We identify several genes that have roles in wing morphogenesis or wing shape variation in *Drosophila* flies, making them promising candidates for future studies. There was little evidence for molecular parallelism in the two species, with only one shared candidate gene, nor for a role of the four known colour pattern loci, except for *optix* in *H. erato*. Thus, we present the first insights into the heritability and genomic basis of within-species wing shape in two *Heliconius* species, adding to a growing body of evidence that polygenic adaptation may underlie many ecologically relevant traits.

## Introduction

Climate change is forcing organisms to ‘move, adapt, or die’. With temperatures rising and land-use changing in the lowlands, shifting to higher elevations might be the only way to flee extinction for many taxa (I.-C. Chen, Hill, Ohlemüller, Roy, & Thomas, 2011). However, the environment changes drastically along mountains, with diverse sets of challenges expected to drive local adaptation (Halbritter, Billeter, Edwards, & Alexander, 2015). Thus, identifying the genomic mechanisms that allow organisms to inhabit wide ranges is key to understanding which taxa are most likely to adapt locally when forced to colonise new, high-elevation, environments. With novel sequencing technologies and the explosion of genomic data, we now have the tools to decipher the genetic basis of ecologically relevant traits in the wild.

Insect flight has many essential functions, including dispersal, courtship, and escaping predators (Dudley, 2002). As such, it is under strong selection to be optimised to suit local environments. Air pressure decreases with elevation, which in turn reduces lift forces required for taking flight, as well as oxygen available for respiration (Hodkinson, 2005). Subtle variation in wing morphology can have big impacts on flight mode and performance (Berwaerts, Van Dyck, & Aerts, 2002). For example, butterflies in tropical rainforests have been found to have rounder wings when they inhabit the understory, both within closely related taxa (Chazot et al., 2014) and across many species (Mena, Kozak, Cárdenas, & Checa, 2020). Elongated wings reduce wing-tip vortices, resulting in more efficient and faster flight, whereas short and wide wings are associated with higher manoeuvrability and lift (Le Roy, Debat, & Llaurens, 2019a). Monarch butterflies have evolved two wing phenotypes; migratory populations have longer wings for long-distance gliding than those that remain year-round in Caribbean islands (Altizer & Davis, 2010). Thus, wing shape represents a trait likely undergoing strong selection across elevations and conferring local adaptation in insects.

Insect wing shape is phylogenetically conserved in many taxa (Houle, Mezey, Galpern, & Carter, 2003; Montejo-Kovacevich et al., 2019; Thomas, Walsh, Wolf, McPheron, & Marden, 2000), but also highly evolvable in the laboratory (Houle et al., 2003), suggesting that wing shape can be heritable. Wing shape tends to have higher heritability than wing size in flies and wasps (Gilchrist & Partridge, 2001; Klingenberg, Debat, & Roff, 2010; Xia, Pannebakker, Groenen, Zwaan, & Bijma, 2020), as size is more prone to life-history trade-offs and condition-dependency. Temperature changes have been shown to trigger plastic responses in a wide range of traits (Deutsch et al., 2008; Merilä and Hendry, 2014; but see Fraimout et al., 2018), including heat tolerance across altitudes in *Heliconius* (Montejo-Kovacevich et al., 2020). Thus, it is important to ascertain the heritability of morphological traits when studying their genomic basis, as plasticity could affect trait values and reduce the power to detect genomic associations. Experimental common-garden designs can help overcome the challenges of studying phenotypic clines in the wild, by providing the same environmental conditions to genotypes of different populations, which allows for the estimation of heritability (de Villemereuil, Gaggiotti, Mouterde, & Till-Bottraud, 2016)

Wing morphology has been mostly studied in *Drosophila*, with developmental pathways identified for *D. melanogaster* (Carreira, Soto, Mensch, & Fanara, 2011; Diaz de la Loza & Thompson, 2017) and many of the candidate genes functionally tested (Pitchers et al., 2019; Ray, Nakata, Henningsson, & Bomphrey, 2016). In other insects, wing dimorphisms have been particularly well-studied. These tend to have simple genetic architectures controlling wingless, or short winged, morphs (B. Li et al., 2020; McCulloch et al., 2019), or are environmentally induced with hormonal regulation, such as in migratory planthoppers (Xu & Zhang, 2017; Zhang, Brisson, & Xu, 2019). However, studies describing the genetic basis of quantitative wing shape variation in organisms other than *Drosophila* are lacking.

Significant advances have been made in understanding the genetic basis of local adaptation in the wild. There are two general strategies to identify loci potentially under selection: (i) forward genetics, where known phenotypic traits are associated to genotypes (via genome-wide association studies or quantitative trait loci with laboratory crosses), and (ii) reverse genetics, where variation in allele frequencies in natural populations is studied to detect signatures of selection across the genome, without any prior knowledge of the phenotypes involved (Fuentes-Pardo & Ruzzante, 2017; Pardo-Diaz, Salazar, & Jiggins, 2015). On the one hand, population genomics, i.e. reverse genetics, excels at detecting genomic regions under selection across environments, but the lack of phenotypic associations can make findings hard to interpret. On the other hand, GWA studies, i.e. forward genetics, require quantifiable traits on a large number of individuals, thus making it difficult to study populations in the wild. A good solution is to study steep clines, where the environment changes continuously over a small space while gene flow is high, and combine both forwards and reverse genetics approaches (Cornetti & Tschirren, 2020; Tigano & Friesen, 2016). Sequencing individuals along such clines allows for sufficient phenotypic variance to measure and associate to genotypes (forward genetics), while maintaining low genetic structure, and additionally test which genomic regions might be undergoing selection by comparing the extremes of the clines (reverse genetics).

The study of aposematic wing pattern colouration in *Heliconius* butterflies is a prime example of this approach. Decades of careful lab crosses described the inheritance of the colour pattern elements and identified the genes controlling them (Martin et al., 2012; Nadeau et al., 2016; Reed et al., 2011; Westerman et al., 2018). In the last decade, whole-genome sequencing in elevationally structured colour-morph hybrid zones has confirmed the role of these loci, which repeatedly diverge across highland and lowland colour pattern races (reverse genetics, Meier et al., 2020; Nadeau et al., 2014). Genome-wide association studies demonstrated that these diverging parts of the genome were associated with specific colour pattern elements. These loci have now been functionally tested in *Heliconius* and other taxa, making important contributions to our understanding of evolution and speciation. Thus, whole-genome sequencing populations across steep environmental clines, while studying phenotype heritability with controlled rearing, is a good approach to disentangle the genomic underpinnings of ecologically relevant traits.

Here we study the genetic basis of wing shape variation in two widespread species of *Heliconius* butterflies across an elevational cline in the Ecuadorian Andes. *Heliconius* inhabiting high altitudes have recently been found to have rounder wings than lowland butterflies, a pattern seen both across species and within species along elevational clines (Montejo-Kovacevich et al., 2019). To estimate the heritability of this potentially adaptive trait, we common-garden reared 71 broods of *H. erato lativitta* and *H. melpomene malleti* from across the cline, yielding 1141 offspring (Fig. 1). We then used forward (GWAS) and reverse (Fst) genetic approaches with whole-genome data of 666 individuals to identify regions associated with quantitative variation in wing aspect ratio and determined which regions diverged between extremes of the cline. This genomic dataset was obtained from a study that developed a new low-cost linked-read sequencing technology, ‘haplotagging’, to examine colour pattern clines in an altitudinally structured hybrid zone (Fig. 1C, Meier et al., 2020). Here we present the first study to examine the heritability and genomic basis of wing shape of two butterfly species in the wild.

**Figure 1.**
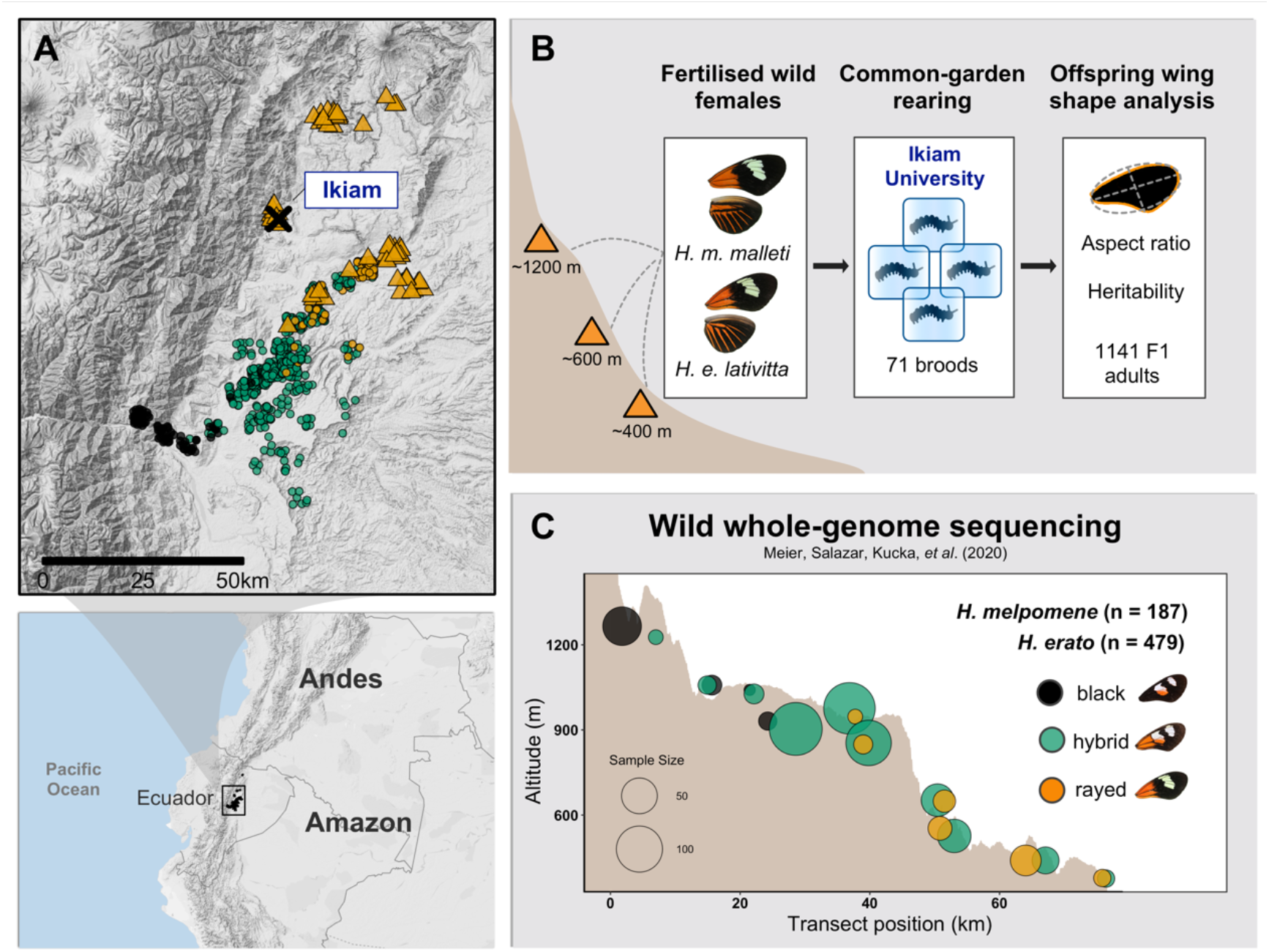
A) Each point represents an individual butterfly collected in the wild (triangles: females used for common garden rearing, circles: individuals whole genome-sequenced with haplotagging (Meier et al., 2020). B) Common-garden rearing protocol. C) Topographic surface of transect across elevations used for whole-genome sequencing. Both species co-occur and have three main colour pattern morphs along this cline: two distinct colour pattern morphs (*H. e notabilis* and *H. m. plesseni*, referred to as “black”, and *H. e lativitta* and *H. m. malletti*, referred to as “rayed”) and within-species hybrids displaying admixed phenotypes (green circles), the most common hybrid phenotype is depicted.

## Methods

### STUDY SYSTEM AND WILD BUTTERFLY COLLECTION

*H. erato* and *H. melpomene* are the two most widespread *Heliconius* species, and have Müllerian aposematic mimicry to advertise their toxicity to predators (Chris D. Jiggins, 2016). They can be found continuously coexisting across elevational clines ranging from sea level up to 1600 m along the Andean mountains. We sampled females of *H. erato* lativitta and *H. melpomene malleti* across the eastern slope of the Ecuadorian Andes for common-garden rearing (Fig. 1B, orange triangles). For the genomic analyses we used a large dataset from a nearby hybrid zone (Meier et al., 2020), where highland subspecies (*H. e. notabilis* and *H. m. plesseni*) meet their respective lowland subspecies (*H. e. lativitta* and *H. m. malleti*) and mate freely, producing a stable intermediate wing pattern population (Fig. 1C). *Heliconius* butterflies were collected with hand nets and precise location recorded. All detached wings were photographed with a DSLR camera with a 100 mm macro lens in standardised conditions, images and full records with data are stored in the EarthCape database (https://*Heliconius*.ecdb.io, Jiggins et al., 2019).

### WING MEASUREMENTS

Wing morphology was analysed with an automated pipeline in the public software Fiji (Schindelin et al., 2012), using custom scripts from Montejo-Kovacevich et al., 2019. These automatically crop, extract the right or left forewing and perform particle analysis on the wing (Fig. 1B). Wing area is estimated for the whole wing in mm^2^. For wing aspect ratio, the particle analysis function obtains the best fitting ellipse of the same area as the wing, extracts the major and minor axis’ lengths, from which aspect ratio is estimated (major axis/minor axis). We only include forewings in this study, as they determine flight speed and mode, whereas hindwings act as an extended surface to support flight and gliding (Le Roy, Debat, & Llaurens, 2019b; Wootton, 2002). Furthermore, hindwings tend to be more damaged in *Heliconius* due to in-flight predation and fragile structure.

### COMMON-GARDEN REARING

Fertilised females of *H. erato lativitta* and *H. melpomene malleti* were caught in the wild with hand nets at elevations ranging from 380 m up to 1600 m (Table S1). Females from all altitudes were simultaneously kept in separate 2×1×3 m cages of purpose-built insectaries at the Universidad Regional Amazónica Ikiam (Fig. 1B. Tena, Ecuador, 615 m). Eggs were collected daily and individuals were reared in separate containers throughout development. Individuals were reared in constant laboratory conditions (21.2 ± 1.1 °C) between 2019-2020, except 10 families from *H. erato* which were reared in common outdoor insectary conditions in 2018. Offspring were individually fed the same host plants, *Passiflora punctata* for *H. erato* and *Passiflora edulis* for *H. melpomene*. Development rates, pupal, and adult mass were recorded for all offspring.

### WHOLE-GENOME DATASET

We used 666 whole-genomes of *H. erato* (n=479) and *H. melpomene* (n=187) from a recent study (Meier et al., 2020), sequenced with ‘haplotagging’. This linked-read sequencing technique retains long-range information via barcoding of DNA molecules before sequencing, which permits megabase-size haplotypes to be reconstructed computationally (Meier et al., 2020). More details on analysis and phasing of molecules can be found in the Supplementary Materials of Meier et al., 2020, and summarised in the Supplementary Materials (Note S1). The resulting dataset used for analyses contained 25.4 million SNP positions for *H. erato* (66.3 SNPs / kbp) and 23.3 million for *H. melpomene* (84.7 SNPs / kbp).

### STATISTICAL ANALYSES

All non-genomic analyses were run in R V2.13 (R Development Core Team 2011) and graphics were generated with the package *ggplot2* (Ginestet, 2011). Packages are specified below and all R scripts are publicly available (Zenodo: TBC). Sequence data from Meier et al. (2020) is deposited at the NCBI Short Read Archive (PRJNA670070).

#### Effects of altitude on wing shape

To test the effects of maternal altitude on wing aspect ratio of common-garden reared offspring, we fitted a linear mixed model that included as fixed effects wing area, sex, development time (days from larva hatching until pupating), and altitude (“high” if the mother was collected above 600m in elevation) with lme4 model fits (Bates, Mächler, Bolker, & Walker, 2015). All continuous fixed effects were standardized to a mean of zero and unit variance to improve model convergence (Zuur, Ieno, Walker, Saveliev, & Smith, 2009). We included family ID as a random effect (intercept) to account for relatedness among offspring in *H. melpomene*. In *H. erato*, we nested family ID within experiment location as an additional random effect, as 10 of the families were reared in common-garden insectary conditions (2018), whereas the rest were reared in laboratory conditions (2019). We performed backward selection of random and fixed effects, in that order, with the package lmerTest (Kuznetsova, Brockhoff, & Christensen, 2017), with likelihood ratio tests and a significance level of α = 0.1 (functions ranova() and drop1() Kuznetsova et al., 2017). When comparing models with different fixed effects we fitted Maximum Likelihood (REML = FALSE), otherwise Restricted Maximum Likelihood models were fitted. Model residuals were checked for homoscedasticity and normality. With the coefficient of determination (R^2^), we estimated the proportion of variance explained by the fixed factors alone (marginal R^2^, R^2^_LMM(m)_) and by both, the fixed effects and the random factors (conditional R^2^, R^2^_LMM(c)_), implemented with the *MuMIn* library (Bartón, 2018; Nakagawa, Johnson, & Schielzeth, 2017).

#### Heritability estimates

To test the heritability of wing shape, we assessed wing aspect ratio variation across individuals from families reared in common-garden conditions. We used all broods with at least three offspring that could be phenotyped. First, we first used an ANOVA approach, with family identity as a factor explaining the variation in aspect ratio. We then estimated narrow-sense heritability (h^2^) with two approaches (more details can be found in Note S2). Firstly, we estimated intra-class correlation coefficient (ICC) or repeatability with a linear mixed model approach; family ID was set as the grouping factor, with a Gaussian distribution and 1000 parametric bootstraps to quantify uncertainty, implemented with the function rptGaussian() in the *rptR* package (Stoffel, Nakagawa, & Schielzeth, 2017). Secondly, we estimated narrow-sense heritability (h^2^) from the slope of mother and mid-offspring wing aspect ratio regressions for those families where the mother’s wings were intact (31/48 broods in *H. erato* and 10/23 in *H. melpomene*). Father’s phenotypes were unknown but their effect should be random, hence unbiased, with respect to the mother’s phenotypes.

#### Genome-wide association mapping of wild wing aspect ratio

All population genomics analyses were performed in ANGSD version 0.933, which uses genotype likelihoods as input to account for genotype uncertainty and a Bayesian framework well-suited for large low-coverage sequencing datasets (Korneliussen, Albrechtsen, & Nielsen, 2014). To account for population structure across the cline, we first calculated admixture proportions with NGSadmix. We used genotype likelihoods computed by STITCH as input and ran NGSadmix on a random subset of 10% of the total sites with a minor allele frequency of at least 0.05, and specified two (k=2) or three (k=3) clusters for *H. erato* and *H. melpomene*, respectively.

To identify the genomic regions potentially controlling quantitative wing shape variation in these two *Heliconius* species, we performed genome-wide association mapping (GWAS, doAsso ANGSD). VCF files from Meier et al. (2020) containing genotype likelihoods were used as input (-vcf-pl). We performed the GWAS with a generalized linear framework and a dosage model (-doAsso 6), which calculates the expected genotype from the input and implements a normal linear regression with the dosages as the predictor variable. Aspect ratio was used as the continuous response variable (-yQuant). Three covariates were incorporated (-cov) to control for sex, wing area, and population structure with the admixture proportions obtained from NGSadmix, as described above. Likelihood ratio test (LRT) statistics are calculated per site (following a chi square distribution with one degree of freedom) under an additive model with logistic regression (-model 1) (Skotte, Korneliussen, & Albrechtsen, 2012).

In GWAS with fewer unlinked SNPs, those with strong associations can sometimes be found in isolation, i.e. without flanking SNPs showing association, and assumed to be in linkage with the causal SNP. However, with large SNP datasets that are not pruned for LD, many are expected to show strong associations when located near the causal site or at random due to noise (Zhou et al., 2020). To partly account for this, we obtained median and minima SNP association p-values for sliding windows of 50 SNP, with a step size of 10 SNP, with the R package ‘WindowScanR’ (Tavares, 2016/2020). When visualising our results, we first use median p-value per window, plotted as −log10(p-value), to regionally smooth the results so that spurious associations get dampened by their flanking high p-value SNPs, while regions with many SNPs with strong associations will be easily identifiable, as their median will remain high. This is analogous to recently developed methods for medical genetics that use penalized moving-window regressions (Bao & Wang, 2017; Begum, Sharker, Sherman, Tseng, & Feingold, 2016; Braz et al., 2019; C. Chen, Steibel, & Tempelman, 2017) or LD clumping (Marees et al., 2018).

We generated a per SNP null distribution of p-values by repeating the genome-wide association analysis 200 times, with randomly permuted phenotypes (aspect ratios) in each run. We then computed median p-values of the same sliding windows across the genome for all 200 permutations. We took the top 1% of windows in the observed GWAS as an initial set of candidate regions. For these we then computed a null distribution of median p-values using the genome-wide permutation p-values for all SNPs in these windows (permutation values shown in white for two regions in Fig. S11). Our final set of outliers only included windows that ranked above the 99^th^ percentile of the window null distribution of p-values, i.e. if the observed median p-value was the lowest or second lowest among the 200 median p-values obtained from permutations. To check for outlier overlaps between species we mapped the *H. melpomene* windows (starts and end positions) to the *H. erato* reference genome using a chainfile from Meier et al. (2020) and the liftover utility (Hinrichs, 2006).

#### Identifying regions diverging between highland and lowland populations

We computed genetic differentiation (Fst) between the highland and lowland subspecies (excluding the mid-elevation hybrids), to identify regions diverging across altitudes and potentially overlapping with wing shape associated regions. With the genotype likelihoods estimated by STITCH, we calculated the site frequency spectrum (SFS) for each population (dosaf, ANGSD). Then, we obtained folded 2D-SFS for both populations combined to use as a prior for the joint allele frequency probabilities with the function realsfs. Fst was calculated per site using the Weir–Cockerham correction (realsfs fst index, ANGSD), and 5kb window averages with 1kb steps were obtained for plotting (following Meier et al. 2020).

#### Candidate gene annotation

Regions associated with wing shape that had at least one 50 SNP window with a median p-value below the threshold of p<0.01 (-log10(p)_median_>2), and ten windows with lower or equal p-values than the lowest 1% of the 200 permutations were investigated further for candidate genes. We obtained the positions of the SNP with the strongest and second strongest association, to get the region with strongest association. We identified one gene if this region was within a gene, or the two closest genes if the region was intergenic or across two genes. We used reference genomes for both species stored in the genome browser Lepbase (Challis, Kumar, Dasmahapatra, Jiggins, & Blaxter, 2016) to identify the genes (*H. erato* demophoon v.1; *H. melpomene H. melpomene* Hmel2.5), and extract the protein sequence. We then searched for similarity in protein sequence databases flybase, uniprotk, and ncbi, and present *Drosophila melanogaster* gene names in the main text and figures. Protein information, E-values, and GO-terms were recorded for each candidate.

## Results

### WING SHAPE PREDICTORS AND HERITABILITY

Wing aspect ratio was phenotyped in 721 *H. erato* offspring individuals from 48 full-sib families, and 419 individuals of *H. melpomene* from 23 full-sib families. Wing aspect ratio varied across families of both species (ANOVA: *H. erato* F_47, 673_ = 3.93, P < 0.0001, *H. melpomene* F_22, 396_ = 11.9, P < 0.0001). Offspring of highland *H. erato* mothers had, on average, rounder wings than those of lowland areas (Fig. 2 B, T-test: *H. erato*: t_46_=-3.5, P<0.001), whereas highland *H. melpomene* offspring were only marginally rounder than lowland families (Fig. 2 D, T-test: t_20_=-2.02, P=0.06), whereas wing size did not differ across broods from different elevations (Fig. S5C). Mean aspect ratio of broods recapitulated those found in wild specimens in a previous study (Fig. S4, Montejo-Kovacevich et al. 2019).

**Figure 2.**
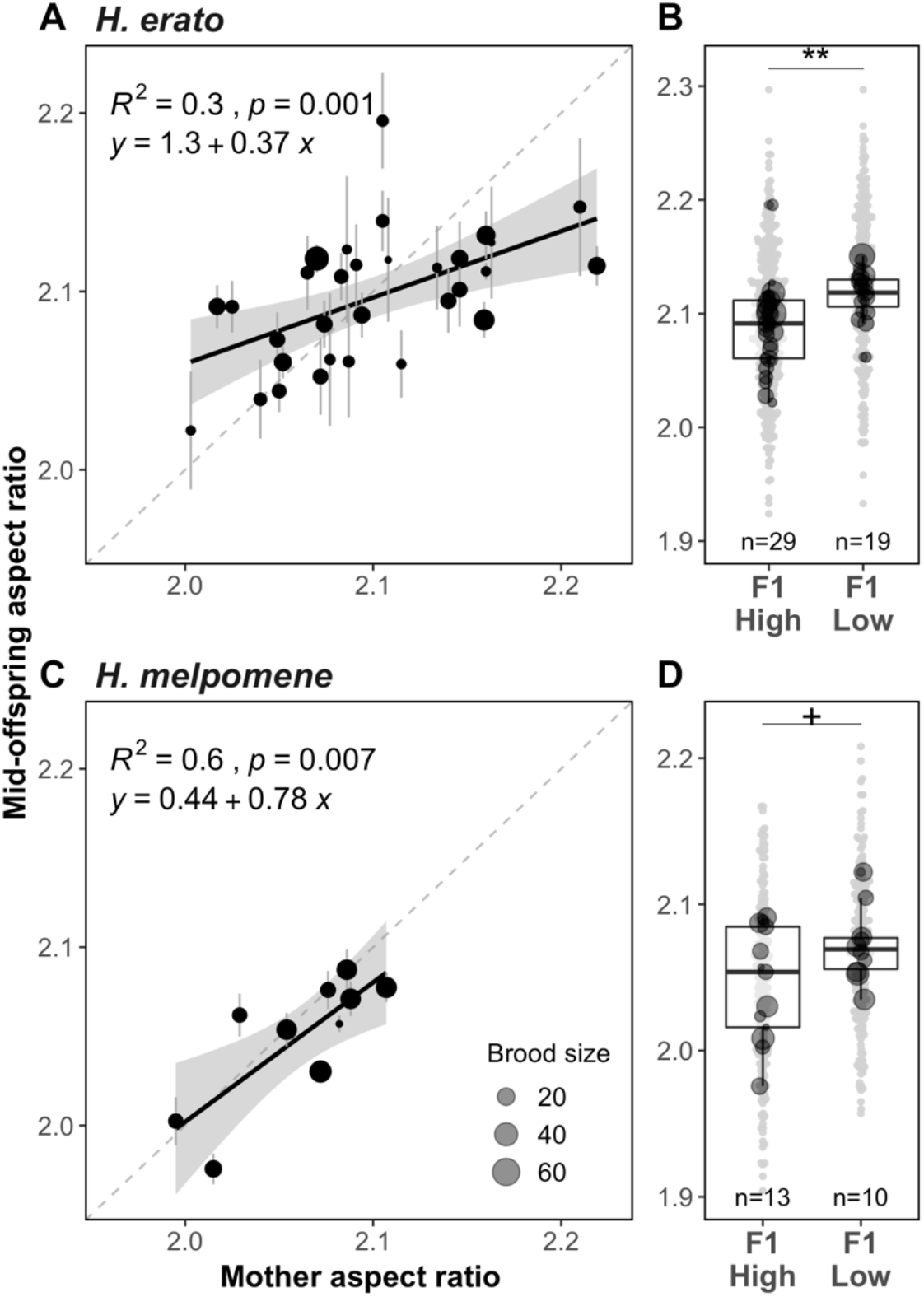
Mother and mid-offspring regression for wing aspect ratio (A,C) and F1 offspring wing aspect ratio with respect to maternal origin across elevations (B,D). Each black point represents mean wing aspect ratio per family and its size is proportional to number of offspring per family. Grey shading around the regression corresponds to 95% confidence intervals of the regression. Stars represent significance levels of two sample t-tests between highland (>600 m) and lowland (<600 m) family means for each species (+<0.075, *< 0.05, **<0.01, ***<0.001).

Altitude, wing area, development time, and sex were significant predictors of wing aspect ratio in common-garden reared individuals of *H. erato*, whereas *H. melpomene* offspring’s wing shape was marginally explained by altitude, with sex and wing size having a stronger effect (Table 1). Of the variation in offspring wing shape, 15% and 38% was explained by family identity while accounting for significant fixed effects in *H. erato* (Repeatability=0.15, S.E.=0.04, P<0.0001) and *H. melpomene*, respectively (Repeatability=0.38, S.E.=0.08, P<0.0001).

**Table 1.** Wing aspect ratio linear mixed model summaries. Fixed effects are scaled and centred. Df, degrees of freedom based on Sattherwaithe’s approximations. *P*, the *P*-values of fixed and random effects. Development time, time in days from larvae hatching to pupating. N, number of individuals with data for all fixed effects. Conditional R^2^ values for models with fixed and random effects (R^2^_LMM(c)_), residual R^2^ values fixed-effects only models (R^2^_LMM(m)_).

| Species | N | Random effects |  |  |  | Fixed effects (scaled) |  |  |  |  |  |
| --- | --- | --- | --- | --- | --- | --- | --- | --- | --- | --- | --- |
| | | Variable | Variance | S.D. | $R^2_{LMM(c)}$ | Variable | Estimate | df | t-value | *P* | $R^2_{LMM(m)}$ |
| *H. erato* | 687 | Mother ID (n=48) | 4.7e-04 | 0.022 | 0.33 | Altitude (low) | 0.029 | 37 | 3.52 | <0.001 | 0.21 |
|  |  | Residual | 2.7e-04 | 0.052 |  | Area | -0.008 | 526 | -3.34 | <0.0001 |  |
|  |  |  |  |  |  | Sex (male) | 0.045 | 658 | 11.02 | <0.0001 |  |
|  |  |  |  |  |  | Development time | -0.008 | 219 | -2.88 | <0.01 |  |
| *H. melpomene* | 419 | Mother ID (n=23) | 1.0e-03 | 0.032 | 0.45 | Altitude (low) | 0.028 | 21.4 | 1.95 | 0.06 | 0.11 |
|  |  | Residual | 1.9e-03 | 0.043 |  | Area | -0.010 | 410.3 | -4.57 | <0.0001 |  |
|  |  |  |  |  |  | Sex (male) | 0.021 | 398.6 | 5.04 | <0.0001 |  |

We obtained the wing aspect ratios for all mothers that retained intact wings in captivity, totalling 31/48 broods of *H. erato*, and for 10/23 of *H. melpomene* broods. Mother and mid-offspring regressions showed strong correlations (Fig. 1 A,C), but heritability was lower in *H. erato* compared to *H. melpomene* (*H. erato*: slope=0.37, R^2^=0.3***; *H. melpomene*: slope=0.79, R^2^=0.6**). Due to the strong sexual wing shape dimorphism in *H. erato*, mother-to-male offspring regressions had a higher intercept (SI Fig. S2A). There was no correlation between mother’s and offspring’s wing size in *H. erato* but a strong correlation in *H. melpomene* (Fig. S3, *H. erato*: slope=0.08, R^2^<0.01 n.s.; *H. melpomene*: slope=0.71, R^2^=0.7**, Fig. S5 A). In contrast, wing size was highly repeatable across broods of *H. erato* (R=0.48, S.E.=0.06), and less so for *H. melpomene* (R=0.15, S.E.=0.05), which could indicate stronger maternal or maternal-environment effects on this trait in *H. erato*.

### GENOME-WIDE ASSOCIATION MAPPING OF ASPECT RATIO VARIATION

#### Subspecies and hybrid phenotypes

Aspect ratio varied slightly across the elevational cline sampled for whole-genome sequencing of both species (Fig. 1 C). Highland subspecies, *H. e. notabilis* and *H. m. plesseni* (black, Fig. 3) were on average rounder than hybrids (green, Fig. 3), and in *H. erato* they were also rounder than lowland subspecies *H. e. lativitta* (orange, Fig. 3 A). Hybrids did not differ in wing aspect ratio compared to their corresponding lowland subspecies, in other words, hybrids were phenotypically more lowland-like (Fig. 3A, Fig. S7, S8). More importantly, the aspect ratio of hybrid individuals encompassed most of the trait variance of the pure subspecies (Fig. S6), increasing our power to detect genomic associations.

**Figure 3.**
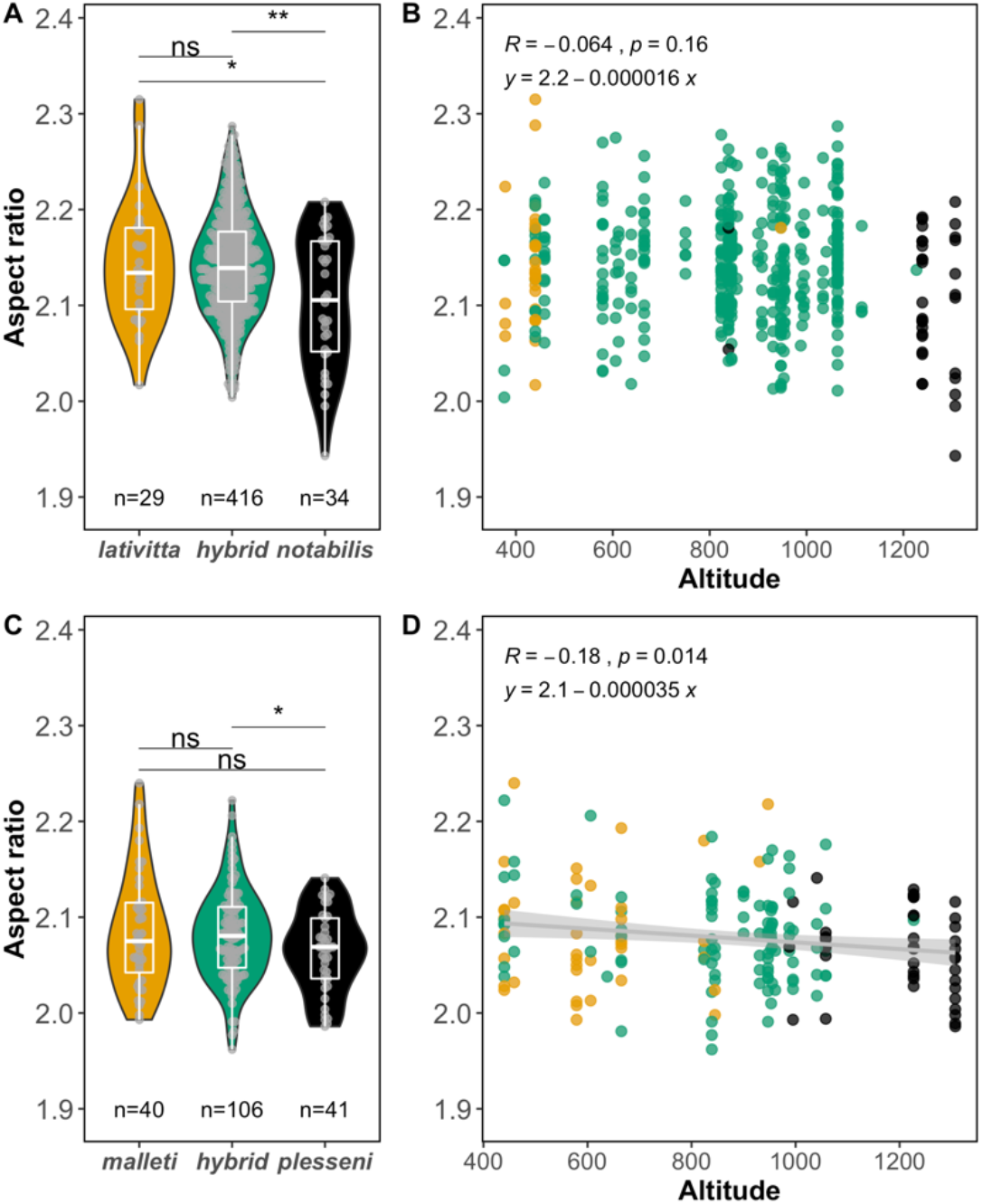
Phenotypes of whole-genome sequenced individuals included in the GWAS (n=666). Wing aspect ratio across subspecies and altitudes of *H. erato* (A, B) and *H. melpomene* (C, D). Stars represent significance levels of two sample t-tests between subspecies of each species (*< 0.05, **<0.01).

#### Association mapping of aspect ratio variation

Genome-wide association mapping revealed a highly polygenic basis for wing shape variation in both species (Fig. 4). We obtained association statistics for 11.3M SNPs (29.4 SNPs/kb) and 10.7M SNPs (38.8 SNPs/kb) for *H. erato* and *H. melpomene*, respectively. Of windows within the 99^th^ percentile association, median p-values, 56% and 39% ranked either first or second lowest among the null distribution obtained from permutations of *H. erato* and *H. melpomene*, respectively (black points in Fig. 4, Fig. S9). Despite a highly polygenic basis, there is evidence that certain genomic regions were strongly associated (Q-Q plots Fig. S10, permutations Fig. S11). We further investigated 28 regions that had 10 adjacent outlier windows supported by permutations and had a median p-value<0.01 (−log10(p)>2, Fig. 4).

**Figure 4.**
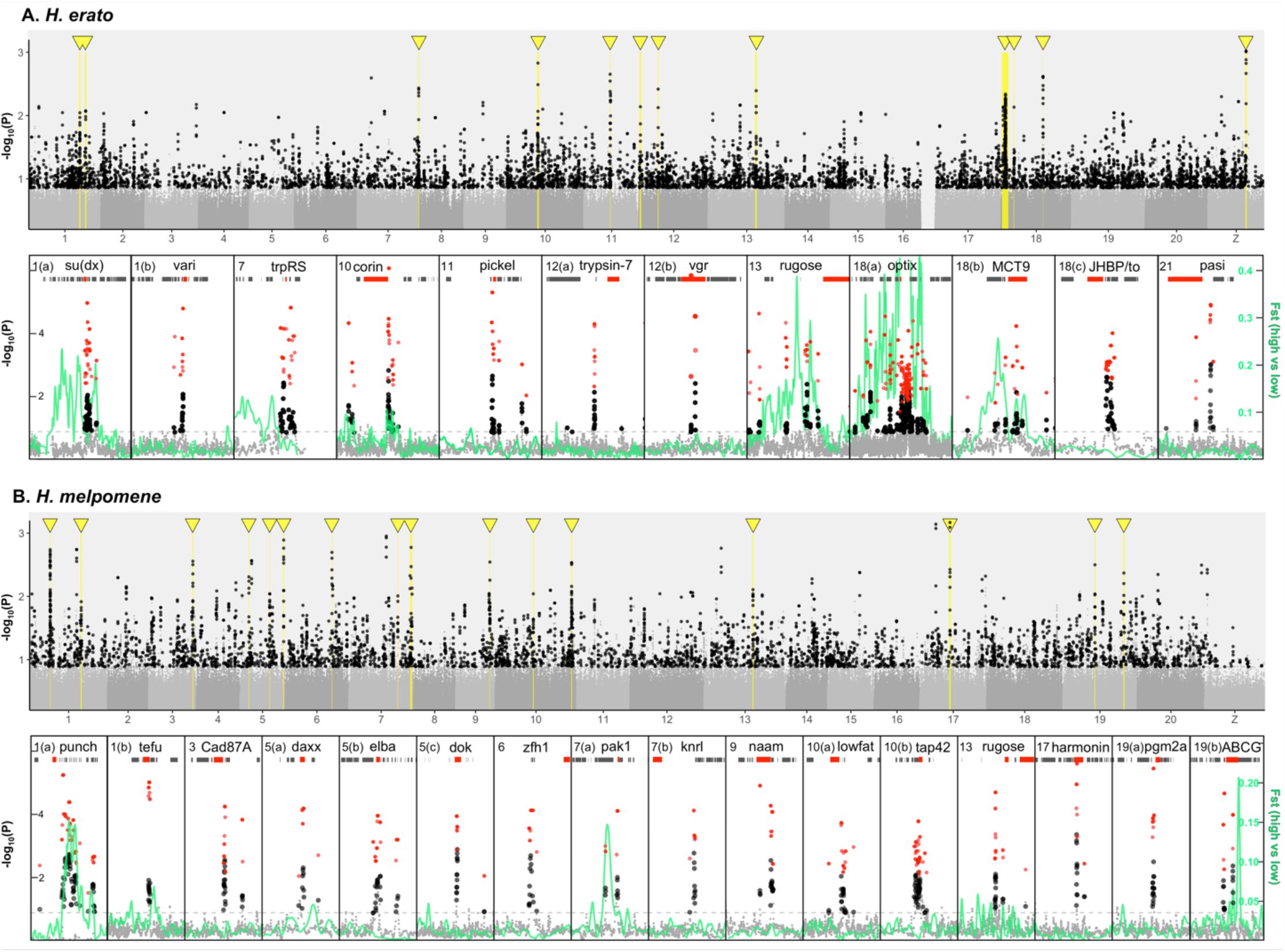
Genome wide association for wing aspect ratio in *H. erato* (A) and *H. melpomene* (B). The windows with the lowest 1% median p-values are above the dotted horizontal grey line, and in black when p-values lower or equal to the top 1% of 200 permutations. Top panels for each species are manhattan plots of genome-wide associations, and bottom are zoomed-in regions of interest. In these regions, minimum p-values per outlier window are shown in red, gene tracks are depicted as grey rectangles, selected genes within or near outlier regions are highlighted in red with gene abbreviations above them, and genetic differentiation between highland and lowland populations along the region in green (Fst).

#### Candidate genes

We identified genes in the 28 regions of interest, within or neighbouring the strongest SNP hits, yielding 23 and 22 candidate genes for *H. erato* and *H. melpomene*, respectively. In 12 out of the 28 regions, SNPs within the region of strongest association were found within genes that could be annotated (Table 2, 3). The remaining outliers were intergenic (n=16) and potentially associated with regulatory variants, and were, on average, 35.8kb from the nearest (n=6) or second nearest gene (n=10, Table 2, 3), i.e. either upstream or downstream. The second nearest gene was presented in the main text if the closest gene was poorly annotated, or if it had a more relevant biological function, e.g. known to affect wing shape or colour patterning in *Heliconius* (all genes in Supplementary Table 1).

**Table 2.** *H. erato* candidate genes for regions of strong association with wing shape variation (n=12). Distance (bp) from gene to outlier window. For those candidate genes that have *Drosophila* mutants leading to wing shape changes we have added references. We highlight (†) candidate or related genes that were identified in Pitchers *et al*. (2019) as significantly affecting multivariate wing shape in *D. melanogaster*. Protein descriptions and relevant biological functions were extracted from Flybase (Thurmond et al., 2019), unless indicated otherwise.

| Chr. | Position outlier window | P.val.min | Gene ID | distance (bp) | Abbrev. | Protein description | Relevant Biological Functions | Mutants in Dmel or GWAS outliers Pitchers(†) |
| --- | --- | --- | --- | --- | --- | --- | --- | --- |
| 1a | Herato0101: 15799088-15801341 | 1.07E-05 | Herato0101.595 | within | su(dx) | E3 ubiquitin-protein ligase, suppressor of deltex | Wing disc dorsal/ventral pattern formation | Vein gaps, rounder wings (Mazaleyrat *et al.*, 2003) |
| 1b | Herato0101: 17600433-17600849 | 1.60E-05 | Herato0101.696 | 1567 | vari | Membrane associated guanylate kinase | Septate junction morphogenesis, open tracheal system |  |
| 7 | Herato0701: 19051639-19051889 | 1.51E-05 | Herato0701.777 | within | trpRS | Tryptophan--trna ligase | Dendrite morphogenesis. Salivary gland. |  |
| 10 | Herato1003: 3435199-3436295 | 8.48E-07 | Herato1003.102 | -3016 | corin | Serine-type endopeptidase | Proteolysis |  |
| 11 | Herato1108: 6393616-6393991 | 4.96E-06 | Herato1108.269 | 2211 | pickel, mega | Claudin transmembrane protein | Septate junction morphogenesis and function. Open tracheal system | Tracheal cell morphogenesis (Behr, Riedel, & Schuh, 2003) |
| 12a | Herato1201: 344400-344590 | 4.61E-05 | Herato1201.11 | 60102 | trypsin-7 | Trypsin-7. Serine-type endopeptidase | Proteolysis |  |
| 12b | Herato1202: 5191025-5191161 | 1.45E-06 | Herato1202.201 | within | vgr | Vitellogenin receptor, yolkless | Oogenesis |  |
| 13 | Herato1301: 14855174-14855376 | 2.32E-05 | Herato1301.557.1 | 68888 | rugose | Neurobeachin | Neuromuscular junction, mushroom body, olfactory learning | Part of epidermal growth factor receptor (*EGFR* †) (Shamloula *et al.*, 2002) |
| 18a | Herato1801: 1382915-1383858 | 1.96E-05 | Herato1801.64 | -131704 | optix | Homeobox containing DNA binding protein | Red colour pattern *Heliconius* |  |
| 18b | Herato1801: 4019579-4019829 | 5.80E-05 | Herato1801.138 | within | MCT9 | Monocarboxylate transporter 9, CG8468 | Monocarboxylic acid transmembrane transporter |  |
| 18c | Herato1805:<br>6957961-6958110 | 9.72E-05 | Herato1805.195 | -2973 | JHBP,<br>takeout | Takeout-like, Juvenile<br>Hormone Binding<br>Protein | Novel circadian<br>output pathway, food<br>status | Reduces male courtship<br>behaviour (Dauwalder,<br>Tsujimoto, Moss, &<br>Mattox, 2002) |
| 21 | Herato2101:<br>11800465-11800678 | 1.22E-05 | Herato2101.392 | -24716 | pasi | CG13288 (Dmel).<br>Uncharacterized<br>tetraspan protein. | Septate Junction -<br>Pasiflora1 (21%<br>resemblance),<br>(Deligiannaki,<br>Casper, Jung, &<br>Gaul, 2015) | Pasiflora1 mutant<br>tracheal barrier<br>breakdown (Deligiannaki<br>et al., 2015) |

**Table 3.**
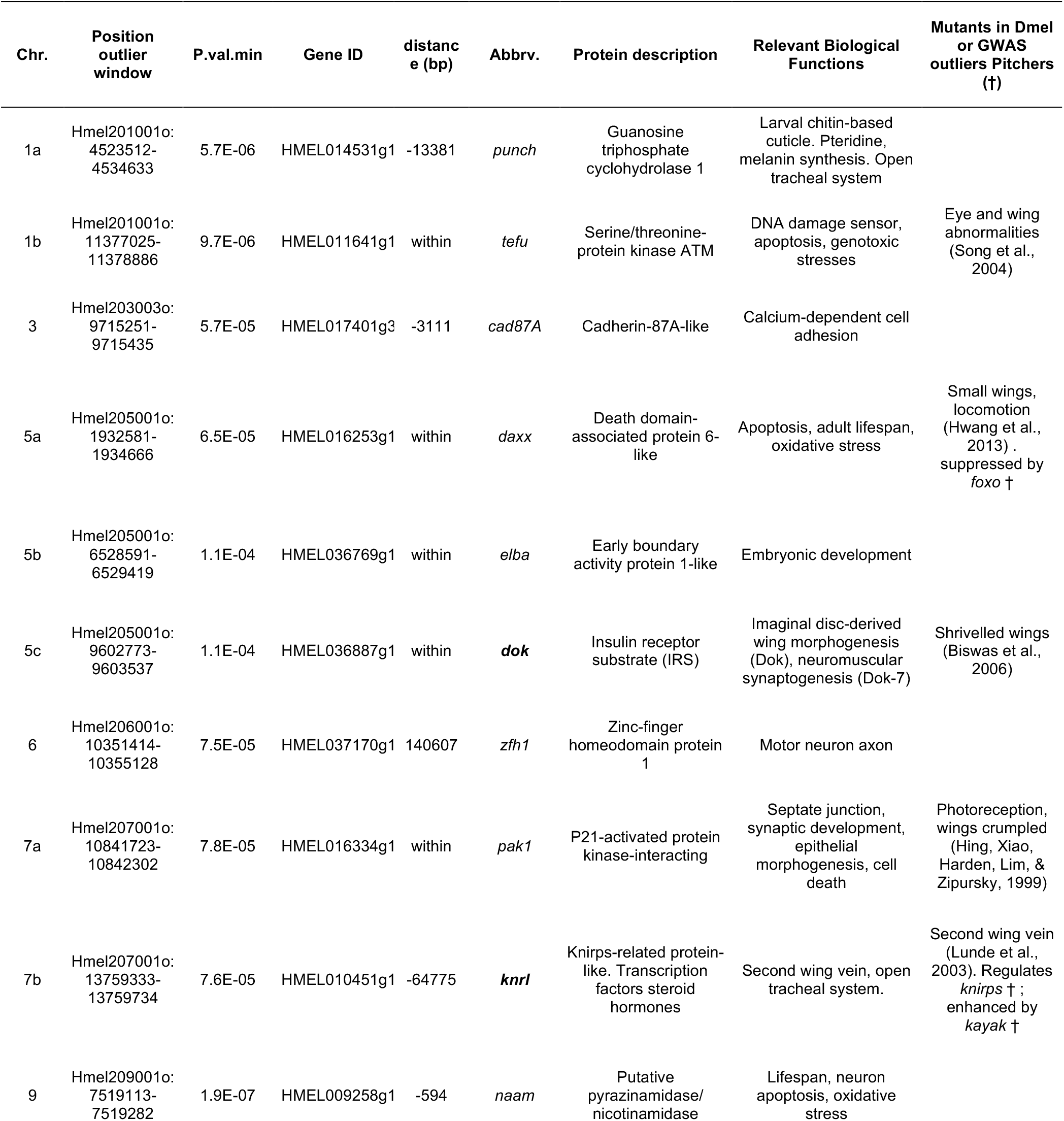
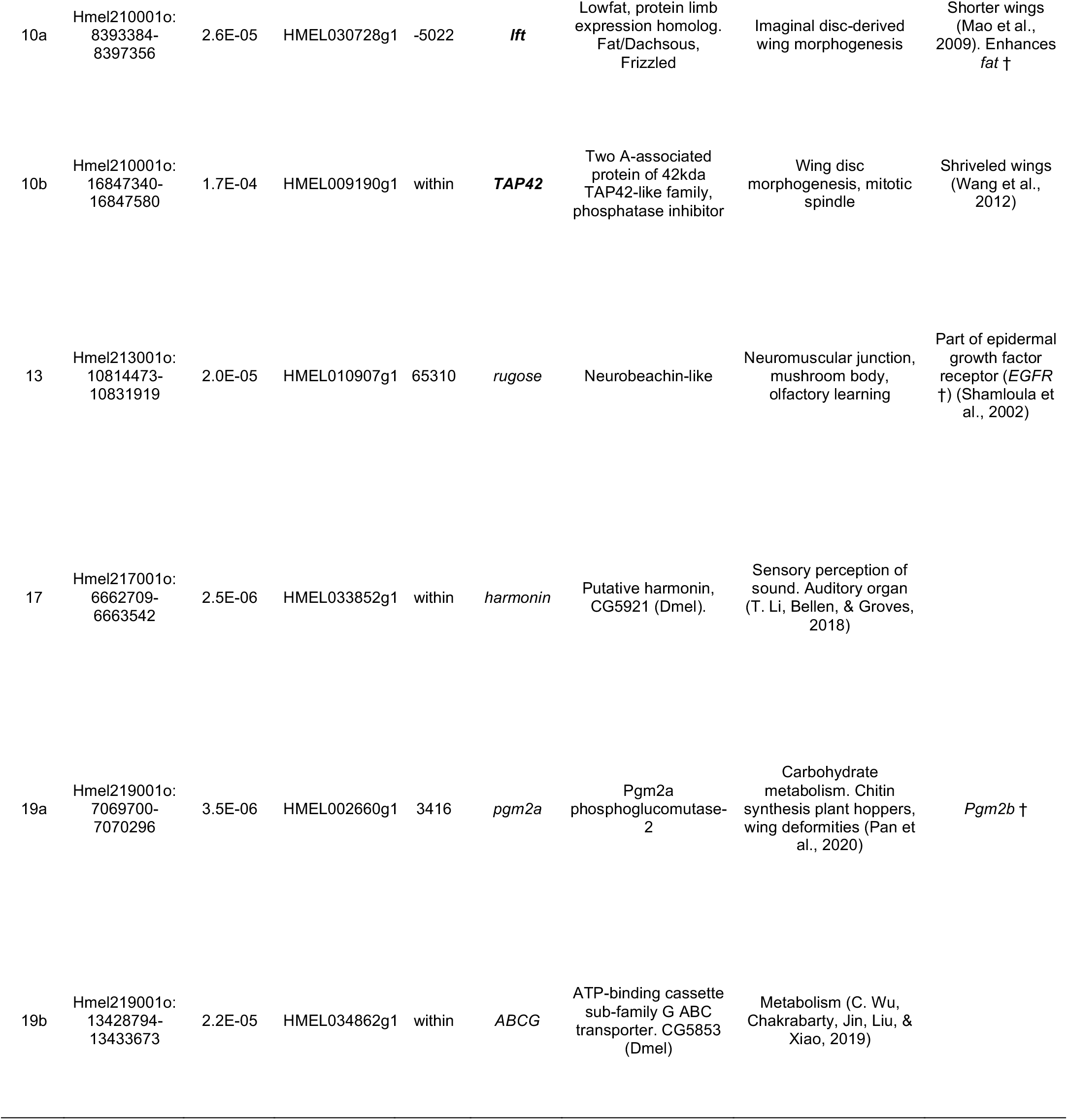
*H. melpomene* candidate genes for regions of strong association with wing shape variation (n=16). Distance (bp) from gene to outlier window. For those candidate genes that have *Drosophila* mutants leading to wing shape changes we have added references. We highlight (†) candidate or related genes that were identified in Pitchers *et al*. (2019) as significantly affecting multivariate wing shape in *D. melanogaster*. Protein descriptions and relevant biological functions were extracted from Flybase (Thurmond et al., 2019), unless indicated otherwise.

Several candidate genes, in both species, encoded proteins previously identified in *Drosophila* as involved in wing morphogenesis. The most relevant and functionally tested candidate genes of wing shape variation were *su(dx)* in *H. erato*, and *dok, knrl, lowfat*, and *tap42* in *H. melpomene* (Fig. 4, Table 2, 3). Tracheal development and septate junction assembly functions were also associated with several candidates (*vari, pickle*, “pasi”, *punch, pak1, knrl*), as well as chitin-based cuticule development (*punch, pgm2a* Pan et al., 2020), pigment transport or synthesis (*MCT9, optix, punch, ABCG*) and oxidative stress responses and regulation of cell apoptotis (*tefu, daxx, pak1, naam*). Some of these candidates, even if not directly involved in wing morphogenesis, have been functionally tested in *Drosophila* and lead to wing shape or wing vein abnormalities (*knrl* Lunde et al., 2003, *pgm2a* Pan et al., 2020, *tefu* Song et al., 2004, *daxx* Hwang et al., 2013). More importantly, 1/23 and 5/22 candidate genes in *H. erato* and *H. melpomene*, respectively, were direct enhancers or suppressors of genes recently identified as being involved with multivariate wing shape variation in *D. melanogaster* (Table 2, 3; Pitchers et al., 2019; Thurmond et al., 2019).

Of the known loci that control colour pattern differences across this cline (Fig. S12, Meier et al., 2020), only one was significantly associated with wing shape variation. The transcription factor controlling presence/absence of red, *optix*, was strongly associated with wing shape in *H. erato*, whereas in *H. melpomene* only a few windows were associated, thus not meeting our criteria for ‘region of interest’ (Fig. 5B, Fig. S11). The outlier region on chromosome 13 was found in close proximity across species, with both SNPs with the lowest p-value mapping nearby the *rugose* gene, which affects neuromuscular junction development, synaptic architecture, brain morphology, and associative learning (Fig. 5A, Table 2, 3. Wise et al., 2015; Zhao et al., 2013). The epidermal growth factor receptor (*Egfr*) regulates *rugose* (Shamloula et al., 2002) and has been associated with wing shape variation in wild *D. melanogaster* (Dworkin, Palsson, & Gibson, 2005) and, recently, in multivariate analyses of wing shape and knockdowns (Pitchers et al., 2019).

**Figure 5.**
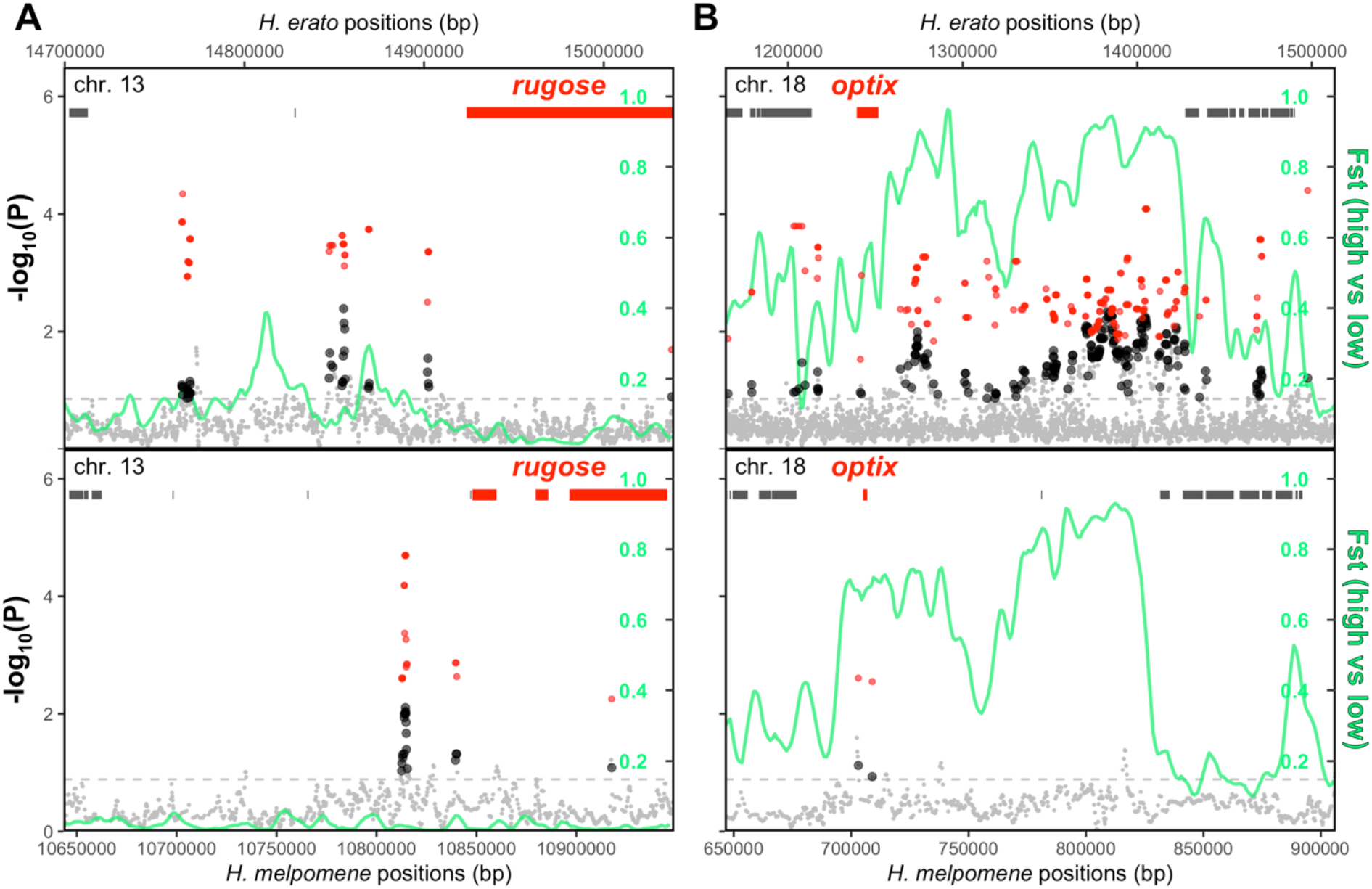
*H. erato* (top) and *H. melpomene* (bottom) region with parallel aspect ratio association at *rugose* (chromosome 13, A), and parallel differentiation (Fst) at the red patterning *optix* locus (chromosome 18, B). The windows with the lowest 1% median p-values are above the dotted horizontal grey line, and in black when above the 99^th^ percentile of 200 permutations. Lowest p-values per outlier window are shown in red, gene tracks are depicted as grey/red rectangles, and genetic differentiation between highland and lowland populations along the region in green (Fst).

### F_ST_ GENOME SCANS

There is little background genomic differentiation between highland and lowland populations of both species (mean F_ST_ in *H. erato*: 0.0261 and in *H. melpomene*: 0.0189, Fig. S12, Meier et al., 2020). Of the 28 regions identified as potentially associated with wing shape variation, ten were found to be in regions of elevated genomic differentiation between highland and lowland populations (F_ST_ green lines, Fig. 4), from low levels of differentiation (e.g. *H. erato* 1(b), Fst_max_=0.08) to high differentiation (e.g. *H. erato* 13, Fst_max_=0.44). The strongest four F_ST_ peaks in this cline are associated with colour patterning (Fig. S9, Meier et al., 2020; Nadeau et al., 2014), but only *optix* (chr. 18) in *H. erato* was also strongly associated with wing shape (Fig. 5B).

## Discussion

Here we combine the power of hybrid zones across steep environmental clines, common garden rearing, and whole-genome sequencing to study the genomic basis of a potentially adaptive trait in the wild. We found that wing aspect ratio is highly correlated between mothers and their offspring in two butterfly species, highly repeatable across common-garden reared offspring families, and correlated with the altitude at which the mother was collected (Fig. 2). With a large dataset comprising 666 whole-genomes sequenced with haplotagging (Meier et al., 2020) and association mapping, we uncover a highly polygenic basis to wing shape, and identify potential candidate genes in regions with many SNPs showing associations (Fig. 4). Furthermore, with a population genetics approach, we find that many of these regions are also diverging between highland and lowland populations, potentially being selected for local adaptation to highland environments.

### WING SHAPE IS HERITABLE

The amount of wing shape variation explained by family across common-garden reared offspring was high for both species (*H. erato*: 21% and *H. melpomene*: 39%), especially when compared to the 74% of variance explained by species identity in a previous comparative study (Montejo-Kovacevich et al., 2019). The resemblance in wing aspect ratio between mothers and their offspring is indicative of a highly heritable trait (Fig. 2 A), although we cannot rule out maternal effects partly driving this pattern. Rearing offspring in common-garden conditions strongly reduces the effects of shared mother-offspring environmental variables, but cannot account for, for example, variation in resources the mothers provide to their eggs. We found, however, that mother wing size did not correlate with offspring wing sizes in *H. erato*, whereas it did in *H. melpomene*. This highlights that wing shape might be less affected by maternal effects compared to other more condition-dependent traits, such as size. Furthermore, strong sexual dimorphism in wing shape present in *H. erato*, was maintained in common-garden reared individuals and both sexes had similar correlations with mother phenotypes, implying a strong genetic component to wing shape variation (Allen, Zwaan, & Brakefield, 2011).

From a local adaptation perspective, we might predict wing shape to be highly heritable. Insects can behaviourally compensate for damaged or abnormal wings through changes in flight and body kinematics (Fernández, Driver, & Hedrick, 2017). Yet, this might incur a fitness cost, as many behaviours, such as courtship and predator escape, are dependent on efficient flight (Le Roy et al., 2019a). Generally, cases of wing shape plasticity are rarer than size plasticity, especially if the two traits are allometrically decoupled, allowing for subtle changes to be selected if advantageous (Carreira et al., 2011; Gilchrist & Partridge, 2001; Strauss, 1990). In *Heliconius*, wings have been found to be rounder at higher elevations, both across and within species that inhabit large ranges (Montejo-Kovacevich et al., 2019). In our study, wing shape differences observed in the wild in *H. erato* and *H. melpomene* were maintained in common-garden reared broods, with individuals from highland mothers having, on average, rounder wings (Fig. 2B, Fig. S4). Together, this supports the hypothesis that subtle changes in wing aspect ratio are highly heritable, and may be involved in local adaptation to altitude.

### CANDIDATES GENES ASSOCIATED WITH WING SHAPE VARIATION IN OTHER INSECTS

We found 5/28 regions mapping to genes involved in the biological process of ‘wing disc development’, which is remarkable given that there are only 419 genes under this category out of 64,000 genes on Flybase (Thurmond et al., 2019), plus one involved in wing vein formation (Lunde et al., 2003). In *H. erato*, the most promising candidate gene was the suppressor of deltex, *su(dx)* (Fig. 4 A, Table 2), an E3 ubiquitin-protein ligase of the Notch signalling pathway (Jennings, Blankley, Baron, Golovanov, & Avis, 2007). *su(dx)* knockouts in *D. melanogaster* result in rounder wings via reduction of longitudinal wing venation (Mazaleyrat et al., 2003), wing margin reduction (Wilkin et al., 2004) or via interactions with other proteins (Djiane et al., 2011). Interestingly, this region has moderate levels of genetic differentiation across high and low elevation populations (Fst_max_=0.24, Fig. 4A, 1(a)), which could be an indication of altitude-associated selection on this candidate gene.

In *H. melpomene*, we found four regions with genes functionally known to be involved in determining wing shape in *Drosophila* (Fig. 4B, Table 3). Mutants of *lowfat* have shorter, rounder wings in *Drosophila* (Hogan, Valentine, Cox, Doyle, & Collier, 2011; Mao, Kucuk, & Irvine, 2009), whereas *dok* mutants have shrivelled wings (Biswas, Stein, & Stanley, 2006). The knirps-related protein (*knrl*) is involved in second wing vein development (Table 3, Lunde et al., 2003), and *Tap42* (Fig. 4B) triggers apoptosis in the developing wing discs (Wang et al., 2012). Interestingly, a recent study found that knockdowns of *fat* and *knirps* affected wing shape in *D. melanogaster*, which regulate two of our candidate genes (Table 3, Pitchers et al., 2019). Here we show a highly polygenic basis to aspect ratio variation in two *Heliconius* species, which contrasts to the simple genetic architecture of wing dimorphisms in other insects (B. Li et al., 2020). The additive effects of many genes could facilitate altitude-associated adaptations, by refining wing shapes to suit the local environment (Wellenreuther & Hansson, 2016). However, aspect ratio may correlate to other wing shape descriptors, thus future studies could focus on multivariate shape variation in these species.

### NOVEL CANDIDATE GENES ASSOCIATED WITH WING SHAPE VARIATION

We found 5/28 regions mapping to genes involved in the biological process of ‘open tracheal system development’, which only has 277 genes on Flybase (Thurmond et al., 2019). Four of these were involved in septate junction assembly function, a category with 83 genes (Thurmond et al., 2019). *varicose*, an *H. erato* candidate gene, is essential to septate junction formation, and *Drosophila* mutants lead to defective tracheal systems and shrivelled wings (Moyer & Jacobs, 2008; V. M. Wu et al., 2007). The veins determine the architecture of the wing, while circulatory and tracheal systems affect the elasticity and physiological functioning of the wing during flight (Pass, 2018). A recent study has shown that butterfly circulatory and tracheal systems are essential for avoiding overheating of the wings by prompting behavioural responses (Tsai et al., 2020). *Heliconius* are generally slow gliding flyers, which display their warning colour patterns to predators, so their wings could rapidly overheat from prolonged exposure to the sun. The potential role of tracheae and wing venation in *Heliconius* flight and thermoregulation remains to be explored.

### CONVERGENCE AND COLOUR PATTERN ASSOCIATION

Despite phenotypic convergence towards rounder wings at high altitude, we found little evidence for molecular parallelism underlying wing shape variation between *H. erato* and *H. melpomene*. One of the 28 regions identified as potentially involved with this trait was found in the regulatory region of the gene rugose in both species (Fig. 5 A). Mutants of *rugose* in *Drosophila* lead to the ‘rough eye phenotype’, similarly to another candidate in *H. melpomene, punch*, which results in cone cell loss and defective vision (Shamloula et al., 2002). *Rugose* mutants also exhibit aberrant associative odour learning, changes in brain morphology, and increased synapses in the larval neuromuscular junction (Volders et al., 2012). A recent study has found *Egfr*, which regulates *rugose*, to affect wing shape in *Drosophila* (Pitchers et al., 2019), making it an interesting candidate for future functional studies. The lack of abundant molecular parallelism in this phenotype is in stark contrast to colour pattern loci in *Heliconius*, which have been repeatedly co-opted or shared via adaptive introgression to create the diversity of mimetic colour morphs we observe across the Neotropics (Fig. S12, Jiggins, 2016; Nadeau et al., 2014).

We found a strong association with wing shape variation in *H. erato* at the *optix* locus, which controls most of the red colour patterning (Bainbridge et al., 2020; Lewis et al., 2019; Meier et al., 2020; Van Belleghem et al., 2017). In *Heliconius*, wing shape has been traditionally studied in the context of mimicry, as similar wing shapes could aid locomotor mimicry (Mérot, Le Poul, Théry, & Joron, 2016; Srygley, 1994, 1999). For example, morphs of *H. numata* that mimic the distantly related genus *Melinea* tend to converge in wing shapes where they co-exist (Jones et al., 2013). Here, wing shape in *H. erato* was more variable between highland and lowland subspecies than in *H. melpomene* (Fig. 3A), which could explain its association with a colour pattern locus (Fig. 5B). It is important to note that, although *H. erato* and *H. melpomene* mimic each other across their range and both have rounder wings in the highlands, the differences in aspect ratio between the two species are larger than those across elevations (Fig. 3D). It therefore seems unlikely that this subtle wing shape variation is primarily caused by mimicry. Furthermore, other species not part of this mimicry ring have been shown to also have rounder wings at high altitudes (Montejo-Kovacevich et al., 2019). Thus, while some mimicry-related wing shape variation may be controlled by *optix*, many other loci are likely involved in shaping wings to suit the local environment and life-history of each species.

### GENETICS BASIS OF AN ECOLOGICALLY RELEVANT TRAIT

A common criticism of population genomics approaches, reverse genetics, that aim to link genotypes and environments is that they often lack phenotypes. Traits directly measured from the wild might be a result of phenotypic plasticity, and are thus rarely used to infer local adaptation. Common-garden rearing can bridge the gap between phenotypes, genotypes, and environment, by providing measurements of heritability and repeatability of a trait across families whose genetic material comes from different extremes of an environmental cline (de Villemereuil et al., 2016). On the other hand, using highly differentiated populations can result in spurious phenotypic associations and makes identifying divergent outliers challenging. Thus, GWAS in the wild should use randomly mating populations with little population structure, while ensuring there is enough phenotypic variation to detect genetic associations with the trait of interest. Hybrid zones, where closely related subspecies or morphs come into contact along an environmental cline, can provide such ideal conditions to carry out GWAS in the wild.

Here we have demonstrated the value of combining these approaches to disentangle the genomic basis of an ecological relevant trait in the wild. We found that wing aspect ratio is highly heritable in two widespread species of *Heliconius* butterflies, and that altitude explains part of the variation in this trait. We have identified several regions potentially shaping wings in *H. erato* and *H. melpomene*, including five candidate genes involved in wing morphogenesis and several identified to be affecting wing shape in recent *Drosophila* studies (Pitchers et al., 2019). We found evidence of molecular parallelism between species at the gene *rugose*, and a strong association of *H. erato* wing shape with a known colour pattern locus, *optix*. Our study adds to a growing body of evidence showing that most quantitative traits conferring local adaptation are highly polygenic (Barghi, Hermisson, & Schlötterer, 2020). Spatial environmental heterogeneity and gene flow are thought to maintain high levels of standing genetic variation (Tigano & Friesen, 2016). This can favour polygenic adaptation, so that incomplete sweeps of many redundant loci can shift traits towards an optimum (Yeaman, 2015). A slow-moving optimum, such range-expansions towards the highlands, should favour polygenic adaptation via small-effect loci, whereas selection for a distant optimum, such a switch in colour pattern mimicry in *Heliconius*, should favour large-effect loci (Barghi et al., 2020). New whole-genome sequencing technologies could foster the study of polygenic adaptation to the environment and shape our understanding of the mode and tempo of evolution in the wild.

## Acknowledgements

We would like to thank all field assistants that have contributed to this study, Narupa Reserve (Jocotoco Foundation), Jatun Satcha reserve, and Universidad Regional Amazónica Ikiam for their support, and Dr. Luca Livraghi for helpful comments. Research permits were granted by the Ministerio del Ambiente, Ecuador (MAE-DNB-CM-2017-0058). G.M.K. was supported by a Natural Environment Research Council Doctoral Training Partnership (NE/L002507/1). Funding was provided to C.B. by the Spanish Agency for International Development Cooperation (AECID, 2018SPE0000400194). Y.F.C. was supported by the European Research Council Starting Grant 639096 “HybridMiX”. This work was supported by the Natural Environment Research Council (NE/R010331/1).

## Author contributions

GMK, PS, FC, CJ, JM, NN designed the study. GMK, PS, SS, KG, CB, NN, performed the experiments and fieldwork. GMK, PS, FC, JM contributed to the analyses. GMK wrote the first draft of the manuscript, all authors contributed to the final version.

## Data availability

All data and scripts are available in the public repository Zenodo (DOI-TBC). Sequence data from Meier et al. (2020) is deposited at the NCBI Short Read Archive (PRJNA670070). All images and data associated to the individuals used for this study are available in the *Heliconius* Earthcape database (https://heliconius.ecdb.io/, Jiggins et al., 2019).

