## Supplementary Information for "Genomics of altitude-associated wing shape in two tropical butterflies"

### *Supporting Information*

**Note S1. Happlotagging datset filtering**

Reads were beadTag demultiplexed, trimmed, placed against their respective references (*H. melpomene*: Hmel2.5 and *H. erato*: helera1_demo), duplicate marked, molecules were identified and initial SNPs called using bcftools call with the multiallelic algorithm (-m). Bam files and initial SNP sets were phased with STITCH, and the resulting variant call files (VCF) were merged and filtered to remove positions with poor information score (INFO_SCORE <= 0.5). Molecular phasing per individual was performed with HAPCUT2 at heterozygous sites. The resulting dataset used for analyses contained 25.4M SNP positions for *H. erato* (66.3 SNPs / kbp) and 23.3M  million for *H. melpomene* (84.7 SNPs / kbp).

**Note S2. Heritability estimates**

There are two ways of estimating narrow-sense heritability ($h^2$) in full-sib designs, where individuals of each group share a mother and a father (Falconer 1995; Caballero 2020). Firstly, resemblance between relatives can be estimated by partitioning phenotypic variance into within-group and between group sources of variance, referred to as intra-class correlation coefficient (ICC) or repeatability. Repeatability (R) is calculated as the variance among group means (group-level variance $V_G$) over the sum of group-level and individual-level (residual) variance $V_R$:

$$R = \frac{V_G}{(V_G + V_R)}$$

This measure of narrow-sense heritability could be inflated, as it includes non-additive genetic effects (dominance or epistasis) and maternal effects, but can be a useful approximation when controlling for across group environmental variance, i.e. in common-garden rearing experiments (Caballero 2020). We estimated within-family wing aspect ratio repeatability (ICC), with a linear mixed model approach. This requires the grouping factor to be specified as a random effect, in this case family ID, with a Gaussian distribution and 1000 parametric bootstraps to quantify uncertainty, implemented with the function rptGaussian() in *rptR* package (Stoffel et al. 2017). By specifying family ID as a random effect, the latter approach estimates the proportion of total wing shape variance accounted for by differences between families. Secondly, narrow-sense heritability ($h^2$) can be estimated with parent-offspring regressions, where mid-parent trait values are regressed against mid-offspring trait values. The slope of such correlation would be considered $h^2$, but can be inflated if trait values are available for only one of the parents. We estimate narrow-sense heritability ($h^2$) by calculating the slopes of mother and mid-offspring regressions, for those families where the mother's wings were intact and phenotyped (31/48 broods in *H. erato* and 10/23 in *H. melpomene*). Father's phenotypes were not available as females were collected fertilised directly from the

**Table S1.** Common garden rearing mothers' locality information. Only broods with more than 3 offspring's wings phenotyped are shown. Unit IDs can be searched in the Earthcape database https://*Heliconius*.ecdb.io, Jiggins et al., 2019.

| Mother ID | Mother unit ID | Species | Batch | Brood size | Locality on Earthcape | Altitude | Latitude | Longitude |
|---|---|---|---|---|---|---|---|---|
| 2 | CAM041354 | *H. e. lativitta* | 18 | 6 | Reventador road | 1312 | -0.01876 | -77.52574 |
| 7 | CAM041833 | *H. e. lativitta* | 18 | 73 | Reventador road 2 | 1348 | -0.004433 | -77.506167 |
| 8 | CAM041393 | *H. e. lativitta* | 18 | 12 | Reventador road 2 | 1348 | -0.004433 | -77.506167 |
| 10 | CAM041429 | *H. e. lativitta* | 18 | 3 | Reventador road 2 | 1348 | -0.004433 | -77.506167 |
| 14 | CAM041473 | *H. e. lativitta* | 18 | 20 | Limoncocha - El Carmen 1 | 458 | -1.09256 | -77.54583 |
| 19 | CAM041510 | *H. e. lativitta* | 18 | 4 | Ikiam Mariposario | 600 | -0.948557 | -77.86605 |
| 20 | CAM041848 | *H. e. lativitta* | 18 | 35 | Road to Shalcana Loma | 429 | -1.057317 | -77.7018 |
| 21 | CAM041568 | *H. e. lativitta* | 18 | 17 | Ikiam Mariposario | 600 | -0.948557 | -77.86605 |
| 22 | CAM041585 | *H. e. lativitta* | 18 | 4 | Reserva Narupa, bridge | 1120 | -0.724668 | -77.767994 |
| 24 | CAM041891 | *H. e. lativitta* | 18 | 49 | Ikiam Mariposario | 600 | -0.948557 | -77.86605 |
| A12 | 19N1981 | *H. m. malleti* | 19-20 | 15 | San Pedro de Arajuno | 405 | -1.09759 | -77.58389 |
| A14 | 19N1694 | *H. e. lativitta* | 19-20 | 8 | San Pedro de Arajuno | 405 | -1.09759 | -77.58389 |
| A15 | 19N2783 | *H. e. lativitta* | 19-20 | 15 | San Pedro de Arajuno | 405 | -1.09759 | -77.58389 |
| A3 | 18N0651 | *H. e. lativitta* | 19-20 | 3 | San Pedro de Arajuno | 405 | -1.09759 | -77.58389 |
| A4 | 19N0270 | *H. e. lativitta* | 19-20 | 22 | San Pedro de Arajuno | 405 | -1.09759 | -77.58389 |
| A5 | 19N1211 | *H. e. lativitta* | 19-20 | 4 | San Pedro de Arajuno | 405 | -1.09759 | -77.58389 |
| E3 | 19N2117 | *H. e. lativitta* | 19-20 | 14 | Reserva Narupa Bridge | 1124 | -0.724668 | -77.767994 |
| H2 | 19N0022 | *H. e. lativitta* | 19-20 | 8 | Challua Yaku Grande | 990 | -0.7185833 | -77.692972 |
| K12 | 19N0326 | *H. e. lativitta* | 19-20 | 14 | Ikiam | 615 | -0.948557 | -77.86605 |
| K17 | 19N0695 | *H. e. lativitta* | 19-20 | 17 | Ikiam | 615 | -0.948557 | -77.86605 |
| K2 | 20N0613 | *H. e. lativitta* | 19-20 | 14 | Ikiam | 615 | -0.948557 | -77.86605 |
| K20 | 19N0981 | *H. m. malleti* | 19-20 | 3 | Ikiam | 615 | -0.948557 | -77.86605 |
| K23 | 20N0614 | *H. e. lativitta* | 19-20 | 5 | Ikiam | 615 | -0.948557 | -77.86605 |
| K24 | 19N2014 | *H. e. lativitta* | 19-20 | 15 | Ikiam | 615 | -0.948557 | -77.86605 |
| K33 | 19N2460 | *H. e. lativitta* | 19-20 | 29 | Ikiam | 615 | -0.948557 | -77.86605 |
| K34 | 19N2463 | *H. e. lativitta* | 19-20 | 6 | Ikiam | 615 | -0.948557 | -77.86605 |
| K7 | 19N0285 | *H. e. lativitta* | 19-20 | 27 | Ikiam | 615 | -0.948557 | -77.86605 |
| L1 | 19N0847 | *H. e. lativitta* | 19-20 | 12 | El Capricho 5.7km | 824 | -1.1878167 | -77.83105 |
| M1 | 20N0615 | *H. e. lativitta* | 19-20 | 30 | La Mina Negra | 1326 | -0.7200278 | -77.747083 |
| M23 | 19N0341 | *H. m. malleti* | 19-20 | 5 | La Mina Negra | 1326 | -0.7200278 | -77.747083 |
| M38 | 19N0528 | *H. m. malleti* | 19-20 | 6 | La Mina Negra | 1326 | -0.7200278 | -77.747083 |
| M39 | 19N0746 | *H. m. malleti* | 19-20 | 25 | La Mina Negra | 1326 | -0.7200278 | -77.747083 |
| M50 | 19N1034 | *H. m. malleti* | 19-20 | 16 | La Mina Negra | 1326 | -0.7200278 | -77.747083 |
| M55 | 19N1467 | *H. m. malleti* | 19-20 | 11 | La Mina Negra | 1326 | -0.7200278 | -77.747083 |
| M59 | 19N1422 | *H. m. malleti* | 19-20 | 15 | La Mina Negra | 1326 | -0.7200278 | -77.747083 |
| N19 | 19N1995 | *H. e. lativitta* | 19-20 | 27 | Finca Narupa bridge | 1150 | -0.72516 | -77.76736 |

| | | | | | | | | |
|---|---|---|---|---|---|---|---|---|
| N23 | 19N1633 | *H. e. lativitta* | 19-20 | 10 | Finca Narupa bridge | 1150 | -0.72516 | -77.76736 |
| N26 | 20N0609 | *H. e. lativitta* | 19-20 | 18 | Finca Narupa bridge | 1150 | -0.72516 | -77.76736 |
| N41 | 20N0616 | *H. m. malleti* | 19-20 | 3 | Finca Narupa bridge | 1150 | -0.72516 | -77.76736 |
| N42 | 20N0617 | *H. m. malleti* | 19-20 | 23 | Finca Narupa bridge | 1150 | -0.72516 | -77.76736 |
| N43 | 20N0618 | *H. m. malleti* | 19-20 | 37 | Finca Narupa bridge | 1150 | -0.72516 | -77.76736 |
| N44 | 19N2510 | *H. e. lativitta* | 19-20 | 39 | Finca Narupa bridge | 1150 | -0.72516 | -77.76736 |
| N45 | 20N0619 | *H. e. lativitta* | 19-20 | 3 | Finca Narupa bridge | 1150 | -0.72516 | -77.76736 |
| N46 | 19N2483 | *H. e. lativitta* | 19-20 | 4 | Finca Narupa bridge | 1150 | -0.72516 | -77.76736 |
| N47 | 19N2784 | *H. e. lativitta* | 19-20 | 8 | Finca Narupa bridge | 1150 | -0.72516 | -77.76736 |
| P1 | 19N0150 | *H. m. malleti* | 19-20 | 13 | Rio Pusuno | 375 | -1.0256944 | -77.60475 |
| P12 | 19N0314 | *H. m. malleti* | 19-20 | 21 | Rio Pusuno | 375 | -1.0256944 | -77.60475 |
| P15 | 19N0007 | *H. e. lativitta* | 19-20 | 13 | Rio Pusuno | 375 | -1.0256944 | -77.60475 |
| P16 | 19N0004 | *H. e. lativitta* | 19-20 | 17 | Rio Pusuno | 375 | -1.0256944 | -77.60475 |
| P19 | 19N0599 | *H. m. malleti* | 19-20 | 25 | Rio Pusuno | 375 | -1.0256944 | -77.60475 |
| P28 | 19N1181 | *H. e. lativitta* | 19-20 | 5 | Rio Pusuno | 375 | -1.0256944 | -77.60475 |
| P29 | 19N1395 | *H. e. lativitta* | 19-20 | 12 | Rio Pusuno | 375 | -1.0256944 | -77.60475 |
| P31 | 19N1413 | *H. e. lativitta* | 19-20 | 7 | Rio Pusuno | 375 | -1.0256944 | -77.60475 |
| P35 | 19N1999 | *H. e. lativitta* | 19-20 | 15 | Rio Pusuno | 375 | -1.0256944 | -77.60475 |
| P39 | 19N1895 | *H. e. lativitta* | 19-20 | 14 | Rio Pusuno | 375 | -1.0256944 | -77.60475 |
| P40 | 19N1646 | *H. m. malleti* | 19-20 | 12 | Rio Pusuno | 375 | -1.0256944 | -77.60475 |
| P41 | 19N2397 | *H. m. malleti* | 19-20 | 37 | Rio Pusuno | 375 | -1.0256944 | -77.60475 |
| P44 | 19N2025 | *H. m. malleti* | 19-20 | 30 | Rio Pusuno | 375 | -1.0256944 | -77.60475 |
| P48 | 19N2785 | *H. m. malleti* | 19-20 | 11 | Rio Pusuno | 375 | -1.0256944 | -77.60475 |
| P49 | 19N2556 | *H. e. lativitta* | 19-20 | 8 | Rio pusuno | 375 | -1.0256944 | -77.60475 |
| P50 | 19N2782 | *H. m. malleti* | 19-20 | 27 | Rio pusuno | 375 | -1.0256944 | -77.60475 |
| T1 | 19N1123 | *H. e. lativitta* | 19-20 | 4 | Shitig | 800 | -0.905718 | -77.873174 |
| T2 | 19N1188 | *H. e. lativitta* | 19-20 | 6 | Shitig | 800 | -0.905718 | -77.873174 |
| U1 | 19N0275 | *H. m. malleti* | 19-20 | 25 | cr. Pununo | 490 | -1.0347778 | -77.656583 |
| V2 | 20N0620 | *H. e. lativitta* | 19-20 | 7 | Venecia | 432 | -0.905718 | -77.873174 |
| W1 | 19N000 | *H. m. malleti* | 19-20 | 15 | Wildsumaco | 1522 | -0.6746389 | -77.608861 |
| W7 | 20N0621 | *H. m. malleti* | 19-20 | 13 | Wildsumaco | 1522 | -0.6746389 | -77.608861 |
| W8 | 19N0280 | *H. m. malleti* | 19-20 | 31 | Wildsumaco | 1522 | -0.6746389 | -77.608861 |
| Y4 | 19N0670 | *H. e. lativitta* | 19-20 | 10 | Apuya 4km | 610 | -1.1155667 | -77.778333 |
| Y6 | 19N0661 | *H. e. lativitta* | 19-20 | 20 | Apuya 4km | 610 | -1.1155667 | -77.778333 |
| Y7 | 19N0320 | *H. e. lativitta* | 19-20 | 8 | Apuya 4km | 610 | -1.1155667 | -77.778333 |

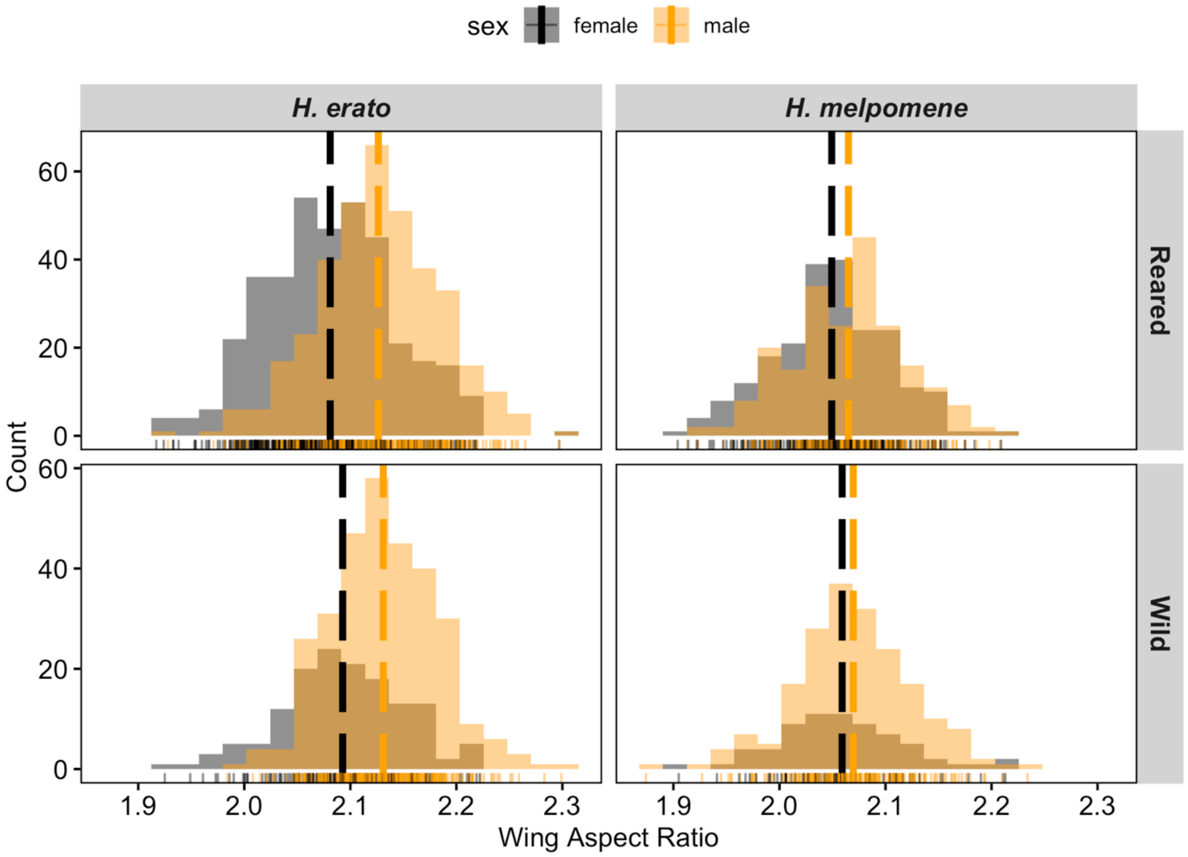


**Figure S1. Sexual dimorphism in *H. erato* and *H. melpomene*.** Wing aspect ratio distribution across sexes in wild and reared individuals included in this study. In the wild, females are captured in smaller numbers due to different diurnal behaviours, hence the lower count for the wild sample. Vertical dotted lines represent means per group.

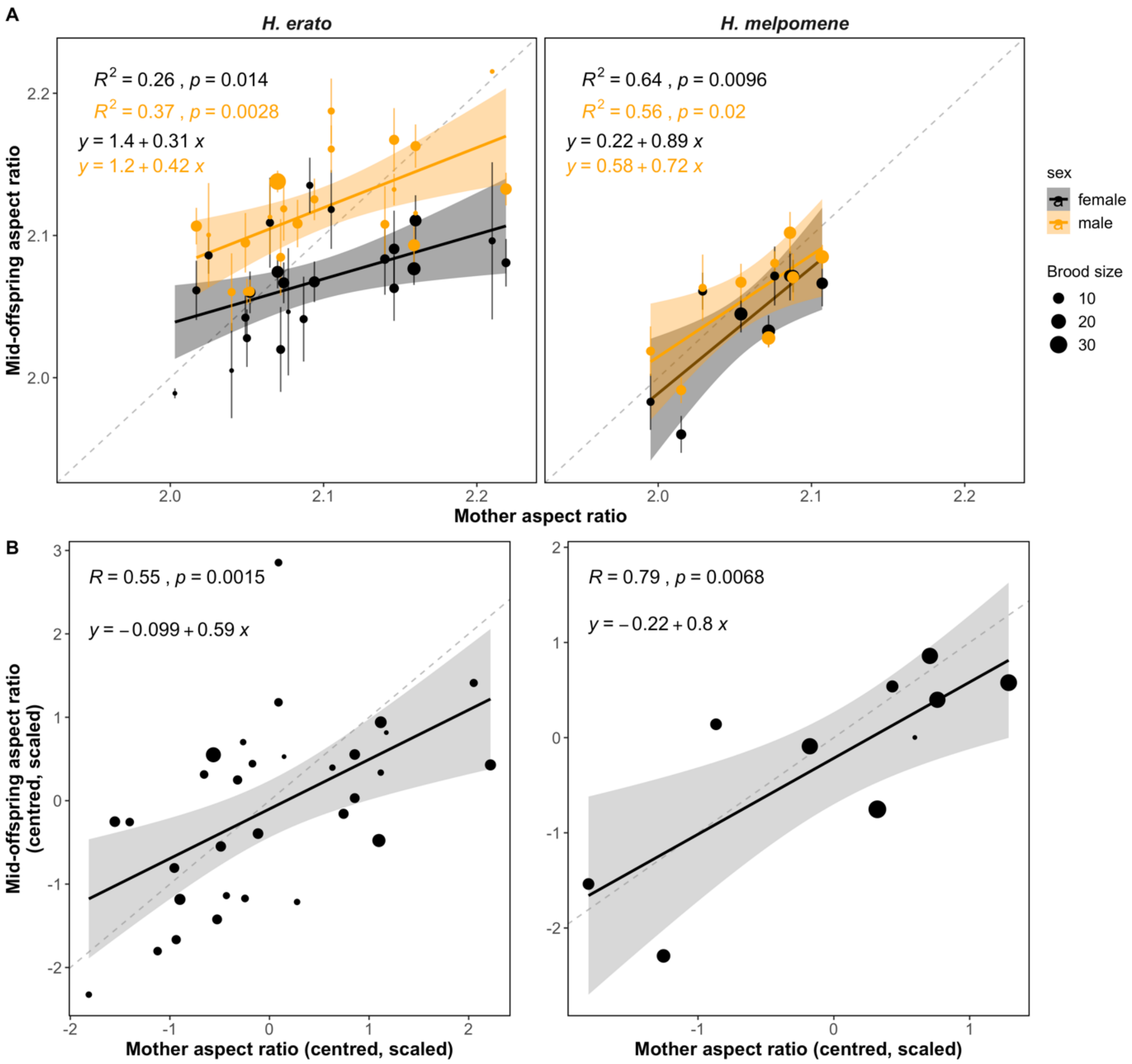


**Figure S2**. Mother and mid-offspring wing shape (aspect ratio) regressions with broods divided into males (orange) and females (A), and with data scaled and centred (B). Left panel: *H. erato*, right: H*. melpomene*. Point size represents number of individuals per brood and vertical lines are standard errors.

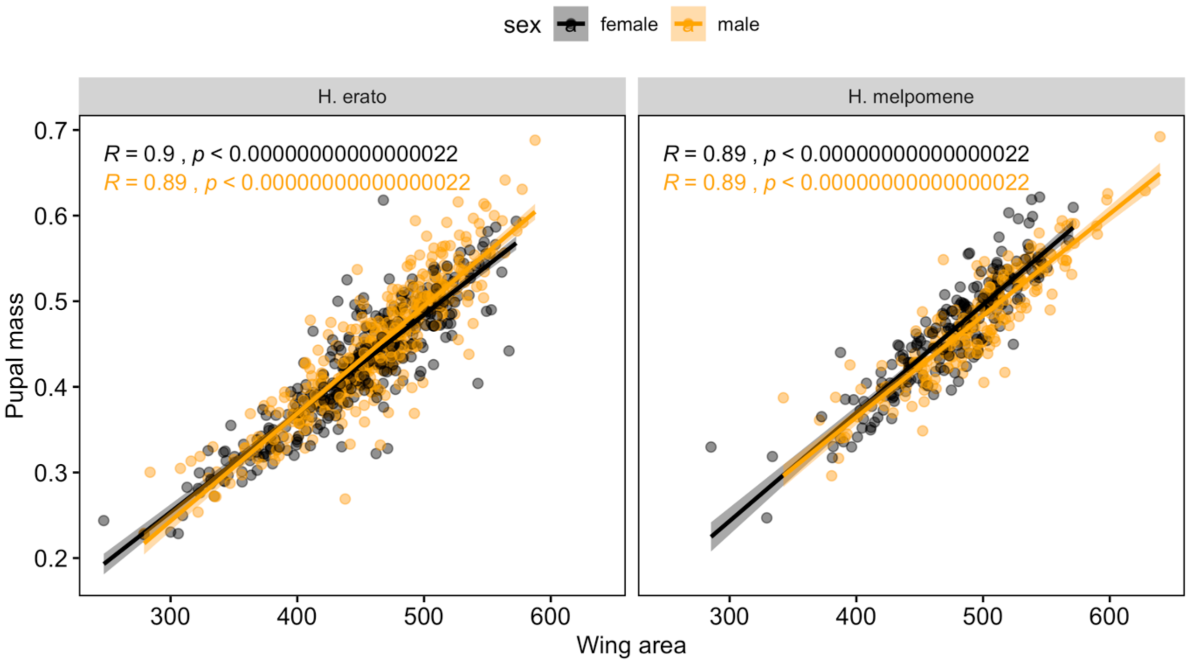


**Figure S3**. Wing area (mm2) of common-garden reared individuals correlates with pupal mass (g) in *H. erato* (right) and *H. melpomene* (left), across sexes. Shading around the regression corresponds to 95% confidence intervals of the regression. Correlation coefficients and p-values of linear regressions are shown.

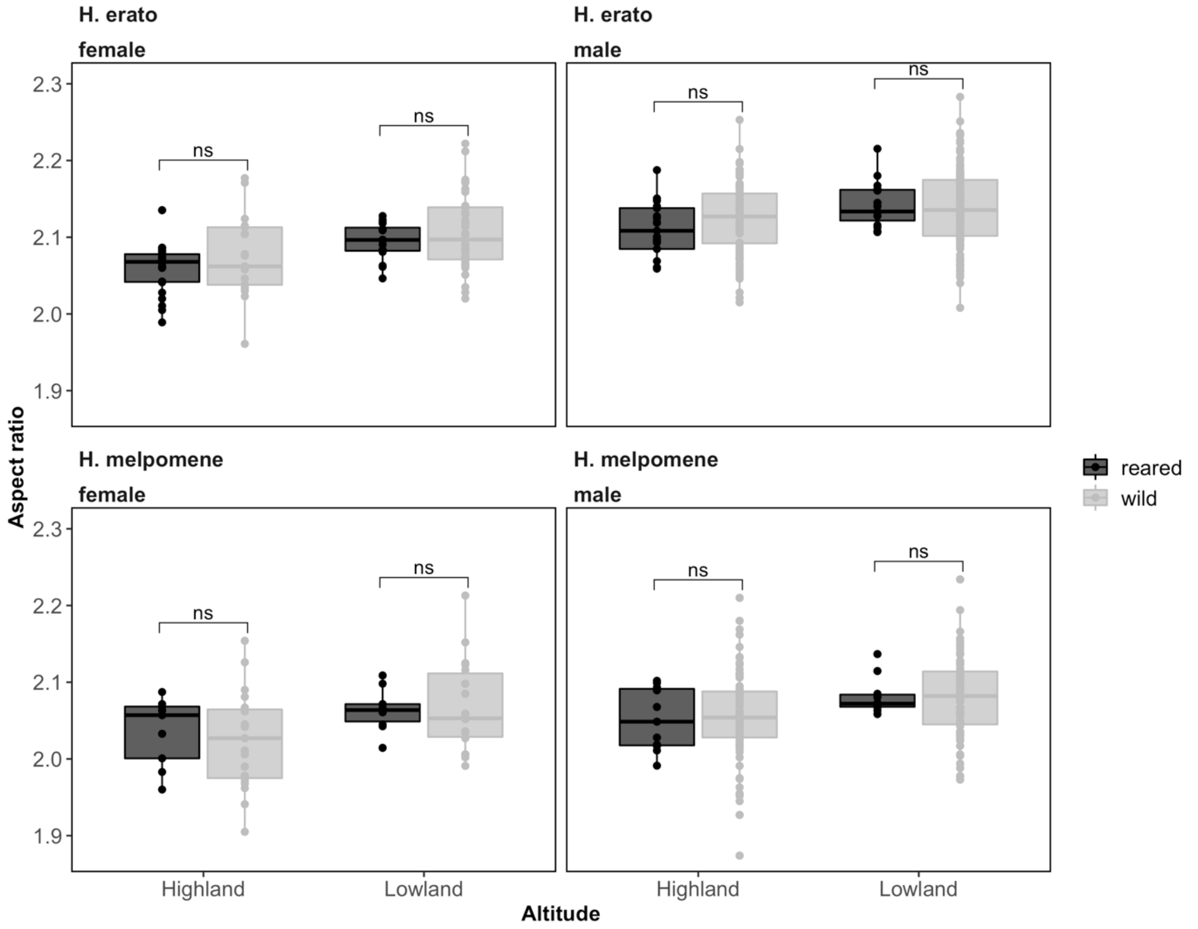


**Figure S4**. Wing aspect ratio from reared broods (black), where each point represents mean aspect ratio per brood, and individuals from the same areas where the brood mothers were collected (grey, from a previous study Montejo-Kovacevich et al. 2019). Two sample t-tests between highland and lowland means for each species and sex were not significant (ns).

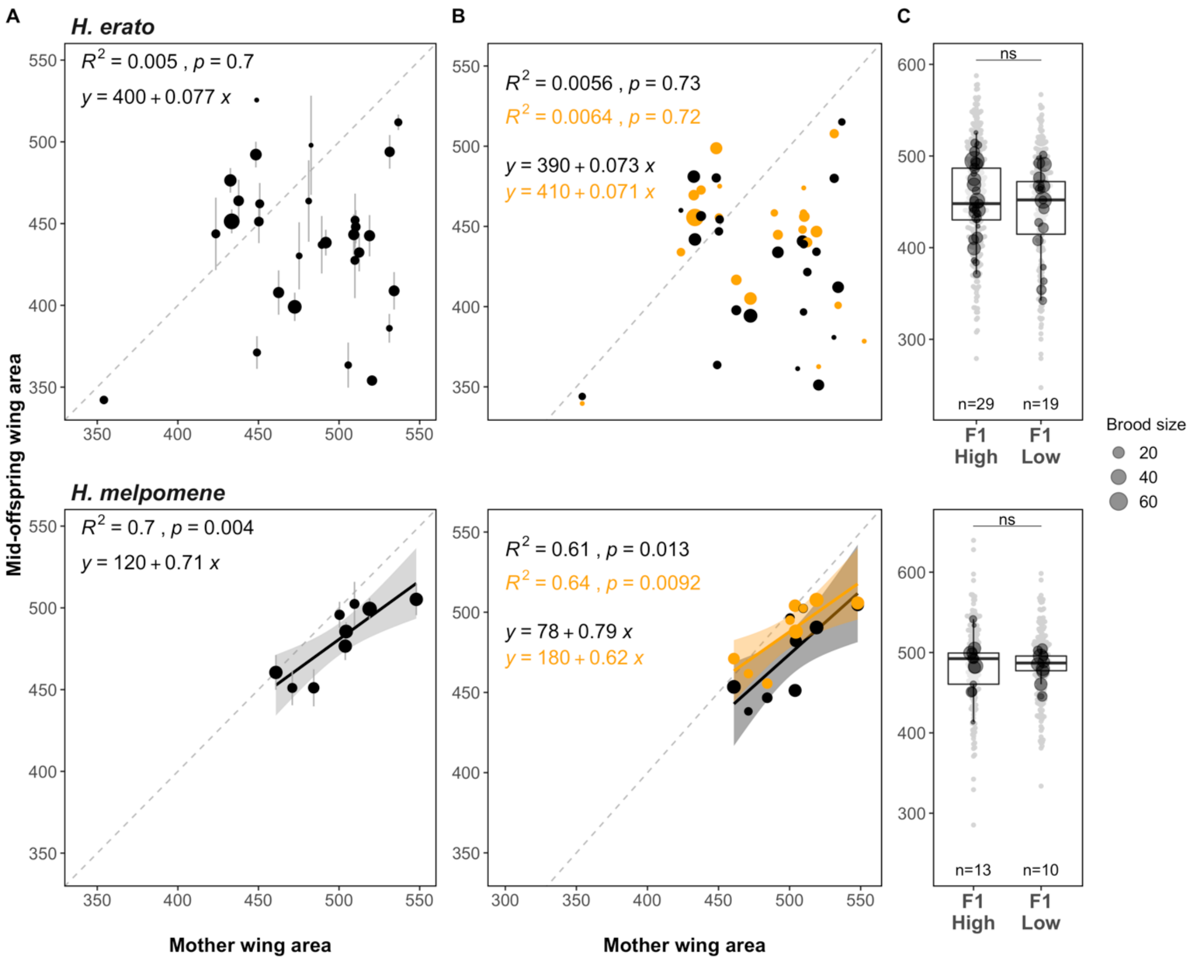


**Figure S5.** Wing area variation across broods and elevations in *H. erato* (top panel) and *H. melpomene*. A) Mother and mid-offspring wing shape (aspect ratio) regressions, B) broods divided into males (orange) and females (black), and C) F1 offspring wing aspect ratio with respect to maternal origin across elevations. Families were classified as high-altitude if the mother was collected above 600 m and low-altitude if below. Wing aspect ratio for all individuals is additionally shown as grey points in the background. Two sample t-tests between highland and lowland family means for each species were not significant (ns).

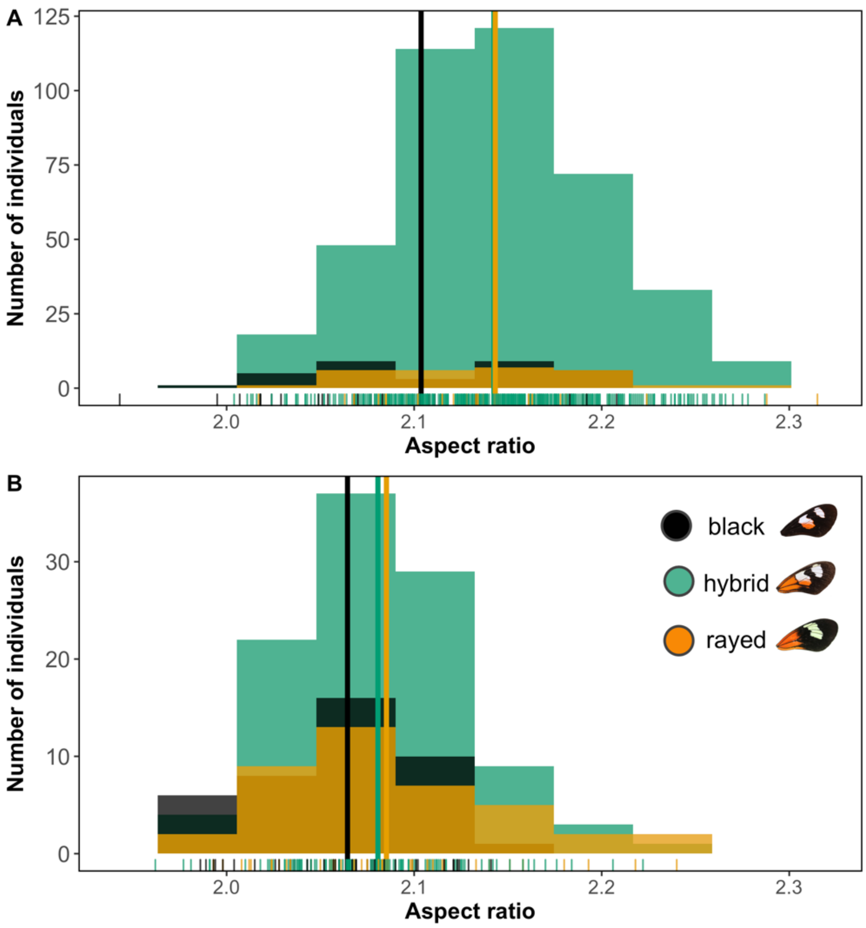


**Figure S6** Wing aspect ratio distribution across subspecies of *H. erato* (A) and *H. melpomene* (B). Both species co-occur and have three main colour pattern morphs along this cline: two distinct colour pattern morphs (*H. e notabilis* and *H. m. plesseni*, referred to as "black", and *H. e lativitta* and *H. m. malletti*, referred to as "rayed") and within-species hybrids displaying admixed phenotypes (green ).

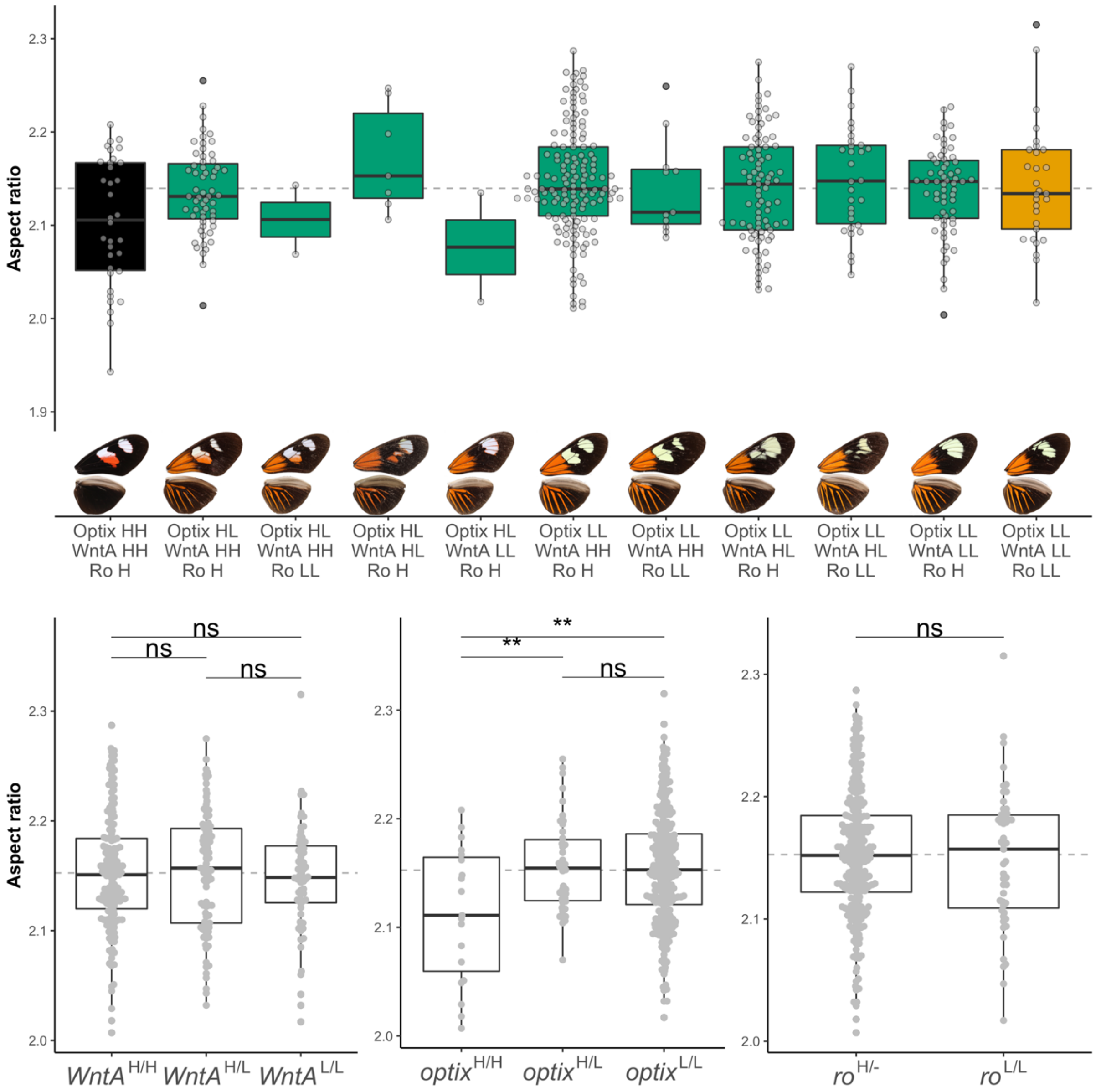


**Figure S7.** Variation in wing aspect ratio across pure subspecies (black= *H. erato notabilis* and orange= *H. erato lativitta*) and intermediate hybrid phenotypes (top) and aspect ratio variation across genotypes (bottom). We divide all individuals into combinations of genotypes at the three most important colour pattern loci for this species: *optix*, *WntA*, and *Ro*, which control red patterns' distribution and presence, number of forewing bands, and forewing band shape, respectively. Highland alleles are represented by "H" and lowland alleles by "L". T-tests of aspect ratio between genotypes are presented (*< 0.05, **<0.01, ***<0.001, ns not significant).

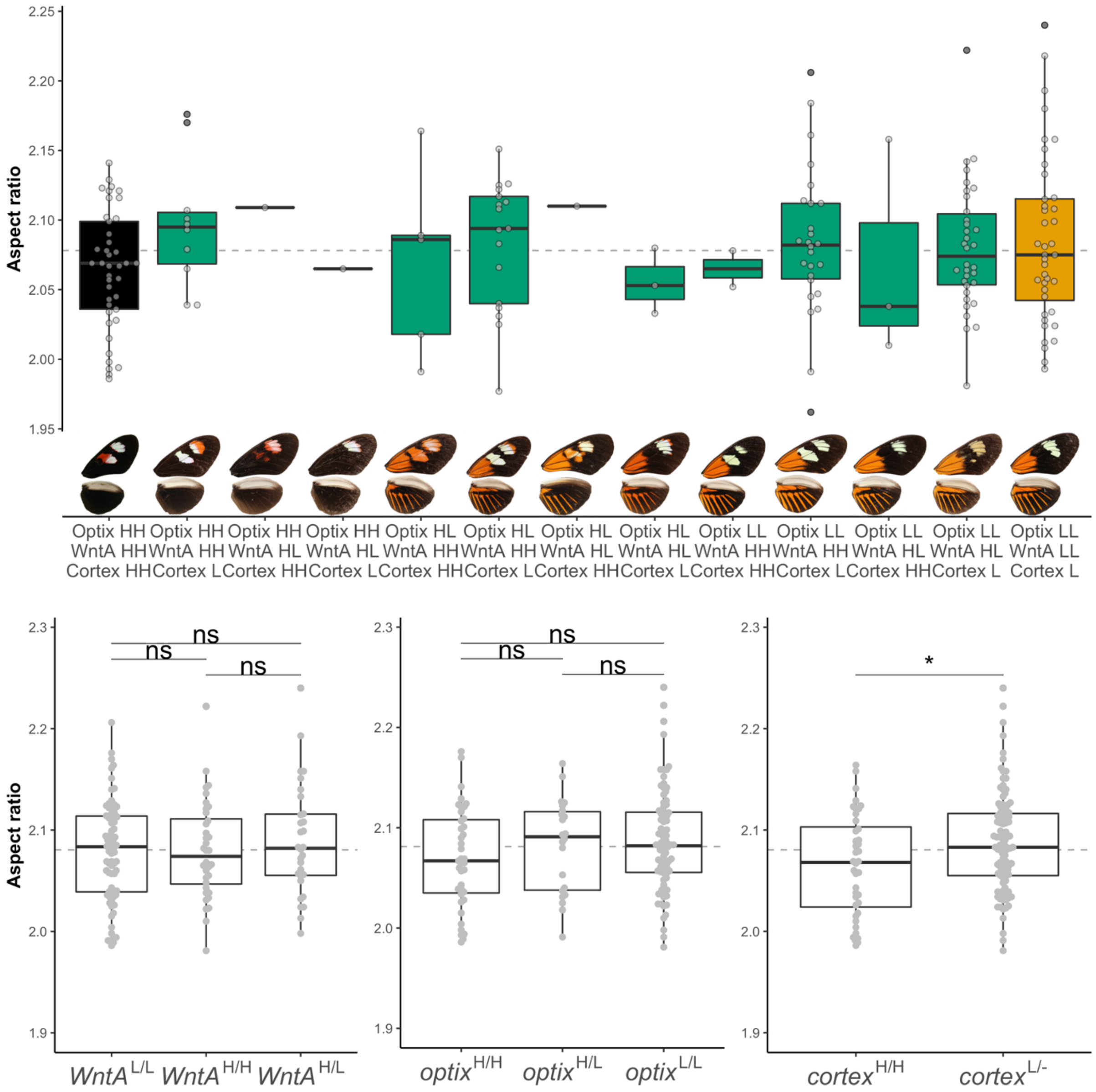


**Figure S8.** Variation in wing aspect ratio across pure subspecies (black= H. erato notabilis and orange= *H. erato lativitta*) and intermediate hybrid phenotypes (top) and aspect ratio variation across genotypes (bottom). We divide all individuals into combinations of genotypes at the three most important colour pattern loci for this species: *optix*, *WntA* and *Cortex*, which control red patterns' distribution and presence, number of forewing bands, and forewing band shape, respectively. Highland alleles are represented by "H" and lowland alleles by "L". T-tests of aspect ratio between genotypes are presented (*< 0.05, **<0.01, ***<0.001, ns not significant).

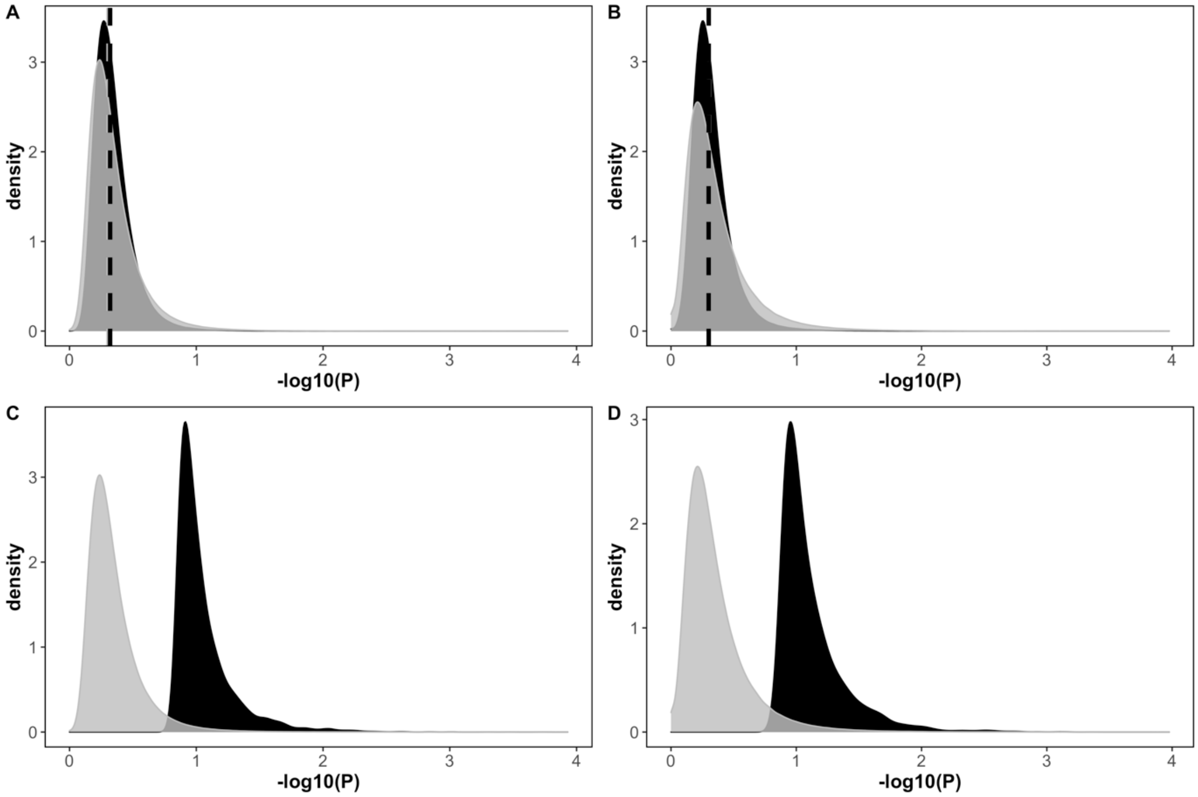


**Fig. S9.** Distribution of p-values (in -log10 scale) under 200 permutations (grey) and observed p-values (black). Top row shows distribution of p-values for all windows (A, B) and bottom row for outlier windows in the observed dataset (i.e. lowest 1% p-values, C, D), for *H. erato* (A, C) and *H. melpomene* (B, D).

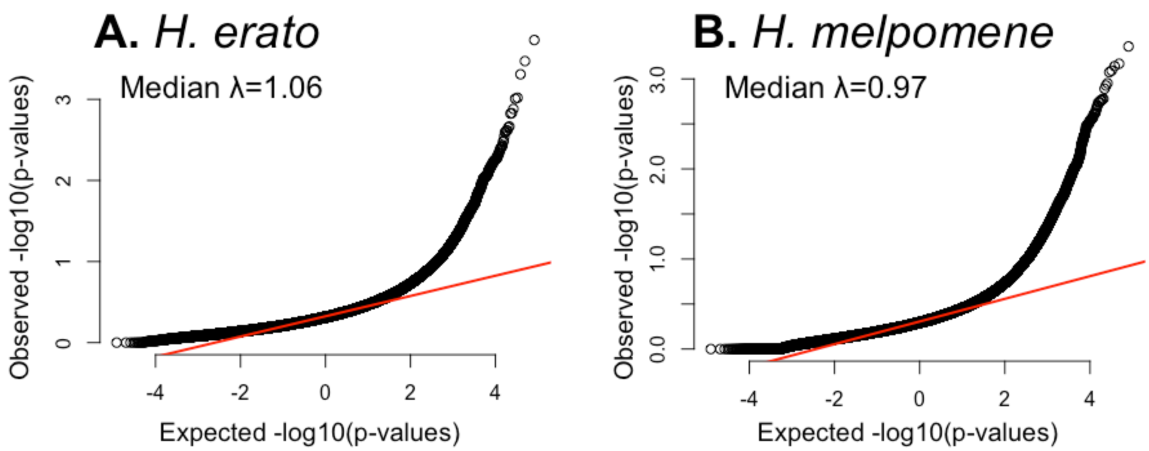


**Figure S10.** QQplot of median p-values per 50SNP window across the genome for *H. erato* (A) and *H. melpomene* (B). Median inflation factor (**λ**) estimated with the function estlambda() from the package GenABEL (Aulchenko et al. 2007).

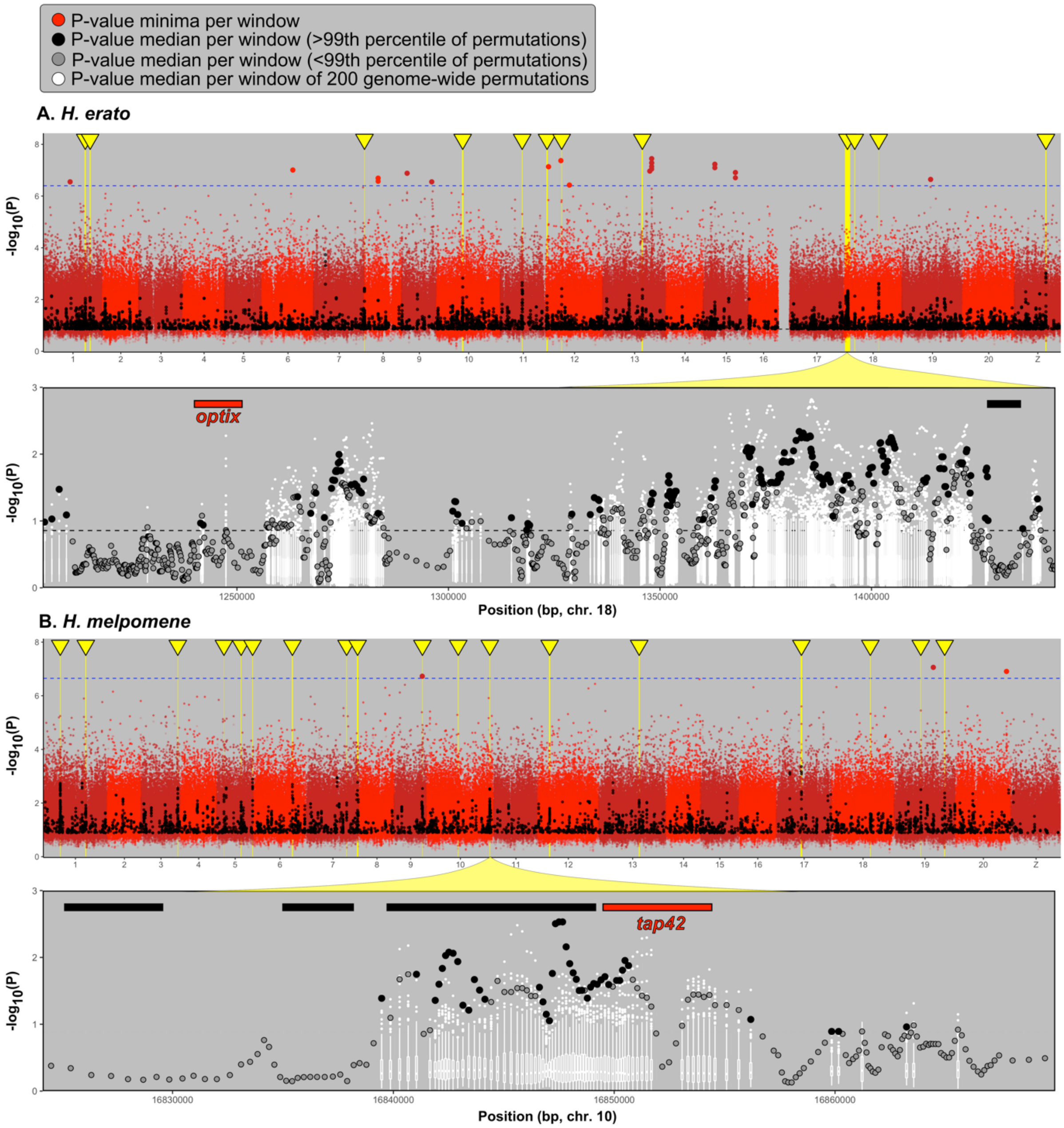


**Figure S11.** Genome-wide association of wing aspect ratio (top panels) and zoomed in regions with median p-values and permutations (bottom panels) in *H. erato* (A) and *H. melpomene* (B). In the genome-wide view, red points represent minima p-value per 50 SNP window (10 SNP step), i.e. the strongest associations, and the blue dotted line shows critical p-value threshold at P<0.05, as obtained from the 95$^{th}$ percentile of minimum genome-wide p-values across 200 permutations. Black points are median p-values per window, for p-values lower or equal to the top 1% of 200 permutations (i.e. observed p-values ranking first or second among permutations). Yellow bars and arrows highlight our regions of interest presented in the main text. The bottom panels are close ups of regions with median p-values only in chromosome 18 (*H. erato*, *optix*) and chromosome 10 (*H. melpomene*, *tap42*). Grey points are median p-values that did not rank above the 99$^{th}$ percentile of permutations, or that were not in the top 1% strongest hits (below the grey dotted line). White boxplots and points represent median p-values for the same windows obtained from 200 genome-wide permutations, where aspect ratio was randomly assigned across individuals.

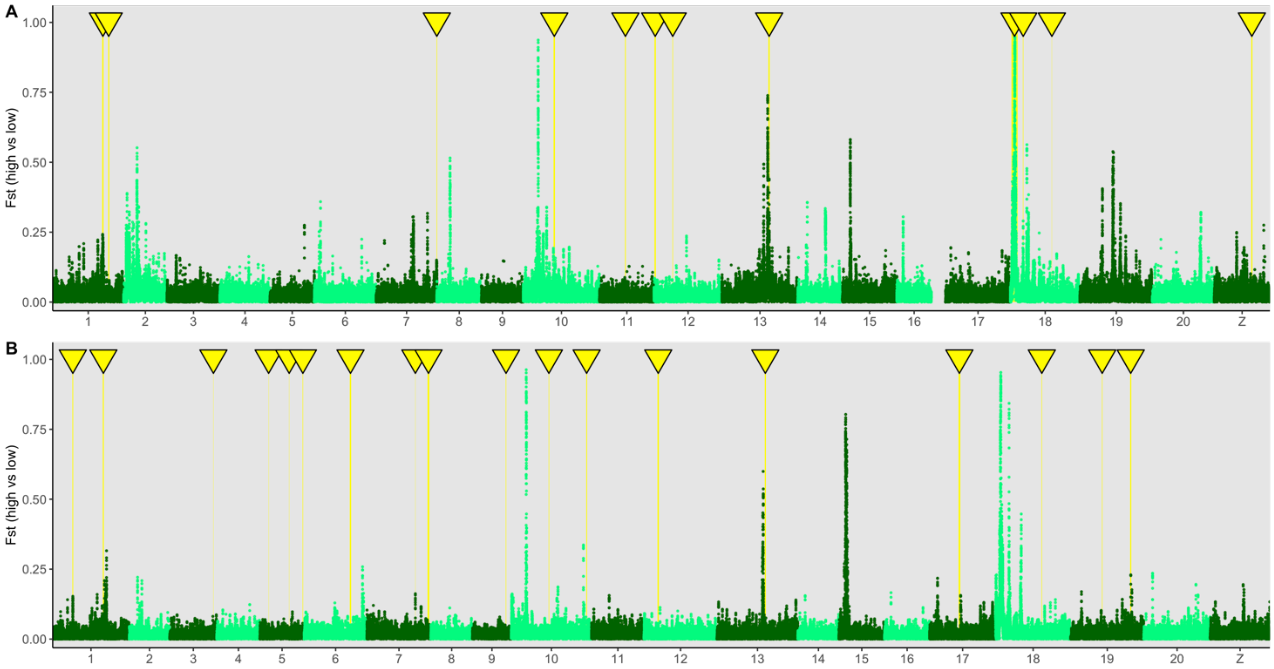


**Figure S12.** Patterns of genetic differentiation (Fst) across the genome of H. erato (A) and H. melpomene (B) high elevation vs low elevation populations. Regions considered outliers in the genome-wide association study for wing shape are highlighted in yellow. The four largest Fst peaks in chromosomes 10, 13, 15, and 18 correspond to known wing pattern loci (described for this cline in Meier et al. 2020).
